# Microbiota-dependent *in vivo* biotransformation, accumulation, and excretion of arsenic from arsenobetaine-rich diet

**DOI:** 10.1101/2024.05.21.594793

**Authors:** Mohana Mukherjee, Lisa Brandenburg, Yuan Dong, Stephanie Pfister, Anika Sidler, Adrien Mestrot, Teresa Chávez Capilla, Siegfried Hapfelmeier

## Abstract

Arsenic (As) is a naturally occurring metalloid that is ubiquitous in the environment. Various chemical forms of As, collectively referred to as “As species”, find their way into the food chain, impacting the health of millions globally. Many of these As species are products of bacterial enzymatic transformations in different environments. Yet, the *in vivo* gut microbial transformation of As and its consequences in the host, especially for less toxic As species like arsenobetaine (AB), remain elusive. Current regulatory assessments regard AB-rich foods, including various seafoods, as safe due to low toxicity and rapid unmodified urinary excretion of the ingested AB. This notion has been challenged by reports of AB metabolism by intestinal bacteria *in vitro* and by more recent evidence of *in vivo* AB metabolism in mice. However, these studies did not definitively establish the causal role of intestinal bacteria in AB transformation *in vivo*. To address this, we employed gnotobiology that allowed us to compare the biotransformation of As from a naturally AB-rich rodent diet in isogenic mice that were either completely germ-free, or colonized with gut microbiota of varying microbial diversity. Additionally, we examined the body distribution, accumulation, and clearance of the ingested As in these mice. Our results confirm the *in vivo* metabolism of AB in the intestine under chronic dietary exposure. The transformation of ingested As was found to be dependent on the presence/absence and complexity of the gut microbiota. Notably, specific toxic As species were absent under germ-free condition. Furthermore, gut microbial colonization was linked to increased As accumulation in the intestinal lumen as well as systemically, along with delayed clearance from the body. These findings emphasize the mammalian gut microbiota as a critical factor in evaluating the safety of AB-accumulating seafoods.

## Introduction

Arsenic (As) is found in nature at variable concentrations and as different chemical species of geological, biological, and anthropogenic origin [1–3]. As is a common contaminant of food and drinking water, and long-term exposure has been linked to various adverse health effects including metabolic, cardiovascular and pulmonary disease, muscle atrophy, neuropathy, paediatric neurocognitive impairment, enteropathy, cancer, and a characteristic precancerous dermatosis of hands and feet (arsenical keratosis) [4–8]. Because of their strong cancer risk association, particularly lung and skin cancers, the two inorganic As species (iAs) arsenate (As^V^) and arsenous acid (As^III^) have been classified as Group 1 carcinogens by the International Agency for Research on Cancer (IARC) [9]. Susceptibility to As-induced diseases has been noted to vary depending on individual genetic makeup, diet, nutritional status, and lifestyle [10]. In addition, the gut microbiota has been identified as a key determinant of the *in vivo* bioavailability, excretion/accumulation kinetics, and toxicity of As [11–16]. For instance, antibiotic-induced depletion of the gut microbiota has been shown to impair *in vivo* As methylation capacity and decrease faecal excretion, along with a concomitant increase in organ accumulation of iAs from drinking water in mice [15, 16]. Other studies in fish and mice revealed that bioaccumulation of As from contaminated diet can be decreased by antibiotic-induced microbial dysbiosis [14, 17]. Gut microbes may affect *in vivo* As metabolism either through direct transformations of ingested As species in the intestinal luminal compartment, or indirectly through modulation of local or systemic host metabolism [14, 16]. Direct intestinal microbial As transformations have been demonstrated *in vitro* utilizing the Simulator of Human Intestinal Microbial Ecosystem (SHIME) culture model [18–20] and *in vitro* microbial fermentation in rodent gut contents [21–24]. Notably, these studies have demonstrated the potential of members of the murine and human gut microbiota to generate the highly toxic As species dimethylmonothioarsinic acid (DMMTAs^V^; IC_50_: 17 µM for human bladder EJ-1 cells for a 24 hour exposure period [25]) and monomethylarsonous acid (MMAs^III^; LC_50_: 18 µM in human UROtsa cells for a 24 hour exposure period [26]) from dimethylarsinic acid (DMAs^V^; IC_50_: > 1 mM for human bladder EJ-1 cells for a 24 hour exposure period [25]) and As^V^ (IC_50_: 85 µM for a 24 hour exposure period in the human bladder carcinoma cell line, T24 [27]), respectively [18, 24]. Prior research has primarily focused on the gut microbiota-mediated transformations of iAs and organic As species such as DMAs^V^ and monomethylarsonic acid (MMAs^V^) that are commonly found in drinking water, soil, and cereals like rice [14, 28, 29]. However, limited information is available on the gut microbial transformations of As from seafood sources [30, 31], particularly arsenobetaine (AB). The organoarsenical AB, primarily found in seafood like fish, crabs, shrimps, and many other marine organisms [32–35] is a predominant source of dietary As exposure for humans and animals worldwide [33, 34, 36–41].

AB-rich foods are considered safe for human consumption based on animal and human studies demonstrating non-toxicity, rapid absorption and urinary excretion, and negligible *in vivo* biotransformation following acute exposure [42–47]. Nevertheless, it has been shown that human large intestinal microbes can degrade AB into other As species, such as DMAs^V^ and As^V^ *in vitro*, challenging the idea of negligible *in vivo* transformation and toxicological inertness of AB in mammals [30]. Although early toxicokinetic human studies concluded that urinary excretion of seafood-derived AB is fast and efficient [44–47], later work indicated that intestinal absorption and urinary excretion of AB from seafood diet may not always be efficient [48–50]. Recent studies have reported gut-microbiota dependent AB accumulation in mice [51] and hypothesized that under chronic exposure scenarios and depending on the specific AB-containing food matrix, a significant portion of AB may reach the large intestinal lumen, where microbial AB degradation can occur, potentially producing arsenic species of health concern [52]. However, these studies did not definitively establish the causal role of intestinal bacteria in these processes. This highlights the need for the systematic investigation of *in vivo* metabolism and accumulation of dietary AB in the mammalian body in dependence of microbial colonization. To differentiate between the individual roles of the host and gut microbes *in vivo*, in the presented study, we employed gnotobiology, a method that allowed us to compare As metabolism in isogenic mice with different microbial colonization statuses. These mice were either completely germ-free (axenic mice, devoid of any live microbes), colonized with a gnotobiotic 12-species bacterial community (known as Oligo Mouse Microbiota-12, Oligo-MM^12^; [53]), or associated with a complex natural mouse microbiota (from conventional specific pathogen-free (SPF) mice obtained from a conventional breeding facility). We compared the *in vivo* biotransformation of As in these mice that were chronically exposed to a naturally AB-rich rodent chow diet as their sole source of As. Furthermore, we compared the body distribution, accumulation, and clearance of the ingested As in these mice.

## Results

### Microbial colonization status influences intestinal luminal accumulation and speciation of As from AB-rich diet

Rodent chow diets used in experimental animal vivaria worldwide are known to commonly contain As concentrations of more than 100 µg kg^-1^, sometimes approaching the European legal limit of inorganic As for animal feed of 2000 µg kg^-1^ [54]. The mice utilized in our study were maintained on a vitamin-fortified autoclavable rodent chow diet (Kliba-Nafag 3307, Granovit CH). The product batch used had a total As concentration of 431 ± 14 (mean ± standard deviation) µg kg^−1^ (**Fig. 1a**). Of this, 92 ± 2 % were identified as AB, 4.0 ± 0.2 % as As^V^, 2.1 ± 0.3 % as DMAs^V^, and 2.1 ± 0.1 % as an unidentified As species (UI00; **Fig. 1b**). Through the exclusion process of quantifying total As in all available diet ingredient samples from the supplier, we could narrow down the likely main As source to fishmeal, suggesting that the high AB content in this diet originates from its natural accumulation in sea fish.

**Figure 1.**
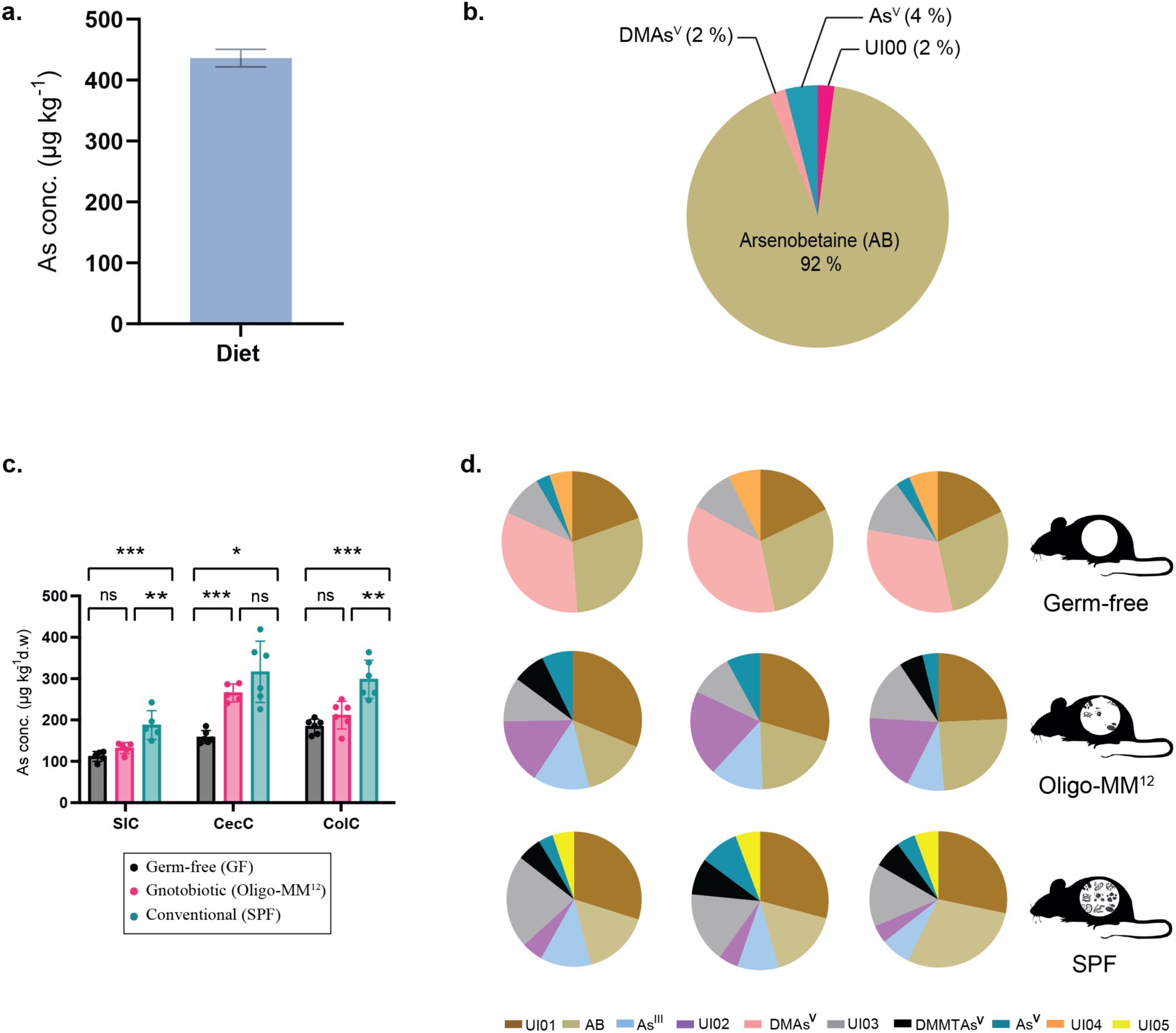
As concentration and speciation in the diet (a and b) and germ-free, Oligo-MM12, and conventional SPF mouse intestinal contents (c and d). (a) Total As concentration and (b) As species composition in the AB-rich chow diet Kliba-Nafag 3307 fed to the mice. Panel (a) shows the mean total As concentration of n = 3 replicates. Error bars indicate the standard deviation. Pie chart in (b) shows mean As species concentrations discovered in n = 3 replicates. AB: Arsenobetaine, As^V^: arsenate, DMAs^V^: dimethylarsinic acid, UI00: Unidentified As species 00. (c) As concentration in the gut luminal contents of C57BL/6J mice during chronic As exposure from an AB-containing diet. SIC: Small intestinal content, CecC: Caecal content, ColC: Colon content. Groupwise mean values are represented by the bar graphs along with individual data points. The error bars represent the standard deviation from the mean value. The normality of each of the data sets was determined with the Shapiro-Wilk test. One-way ANOVA was performed to identify significant differences between the mean values of each of the mouse groups. (SIC: ***, adjusted p-value = 0.0005; **, adjusted p-value = 0.0057; ns, adjusted p-value = 0.3756; Ordinary One-way ANOVA. CecC: ***, adjusted p-value = 0.0001; *, adjusted p-value = 0.0101; ns, adjusted p-value = 0.3722; Brown-Forsythe and Welch ANOVA. ColC: ***, adjusted p-value = 0.0001; **, adjusted p-value = 0.0016; ns, adjusted p-value = 0.3875; Ordinary One-way ANOVA). The SPF mice show significantly higher As concentrations than the germ-free mice in all three compartments, while, the As concentrations in the Oligo-MM12 mice are much more comparable to the germ-free mice except for the caecum content. (d) As species identified in the caecal contents of germ-free (upper panel), Oligo-MM12 (middle panel), and SPF mice (lower panel) in triplicates. Each pie chart is representative of an individual mouse caecum As metabolite profile. Limit of detection (LOD) = 0.095µg kg^-1^. UI01: Unidentified species 01, AB: Arsenobetaine, As^III^: arsenous acid, UI02: Unidentified species 02, DMAs^V^: dimethylarsinic acid, UI03: Unidentified species 03, DMMTAs^V^: dimethylmonothioarsinic acid, UI04: Unidentified species 04, UI05: Unidentified species 05.

We first determined whether microbial colonization influenced the gut luminal accumulation of As from the AB-rich diet. **Fig. 1c** summarizes the total As concentrations in small intestinal (SIC), caecal (CecC), and colon contents (ColC) of germ-free, Oligo-MM^12^, and conventional (SPF) mice. SPF mice, harbouring the most diverse microbiota, showed the highest As concentrations in all intestinal segments and in the faeces (see **Supplementary material: Section III Supplementary Fig. S9**), followed by the Oligo-MM^12^ and germ-free mice. While SIC and ColC As concentrations of 132 ± 13 µg kg^−1^ and 212 ± 34 µg kg^−1^ in Oligo-MM^12^ mice were only slightly higher than in germ-free mice (SIC: 112 ± 12 µg kg^−1^; ColC: 185 ± 18 µg kg^−1^), their CecC and faecal (see **Supplementary material: Section III Supplementary Fig. S9**) As levels (266 ± 21 µg kg^-1^ and 311 ± 23 µg kg^−1^, respectively) approached those in SPF mice (317 ± 74 µg kg^-1^ and 350 ± 56 µg kg^−1^, respectively) and were significantly higher than in germ-free mice (159 ± 16 µg kg^-1^ and 206 ± 15 µg kg^−1^, respectively). Hence, association with a gut microbiota increased large intestinal luminal accumulation of As from an AB-rich diet, reminiscent of prior reports of microbiota-dependent faecal accumulation of As from As^V^-containing drinking water [15, 16].

To examine the *in vivo* gut microbial and host As transformations associated with these toxicokinetic differences (see **Fig. 1c**), caecal contents from a sub-cohort of these germ-free, Oligo-MM^12^, and SPF mice (n = 3) were further analysed for As speciation. **Fig. 1d** shows the As species compositions in the caecal contents of these mice. Preliminary As species identification was done by comparing peak retention times (R_t_) of the sample chromatograms with those of As species standards in an ion-pairing chromatography. Further confirmation or characterization of some As species were attained through various approaches (**Supplementary material: Section II**). AB and As^V^ were seen in all three mouse groups. Although receiving the same diet, the germ-free intestinal As metabolite profile was markedly different from that of Oligo-MM^12^ and SPF mice. The host-derived caecal As species metabolome of the germ-free mice was predominated by DMAs^V^ followed by AB and the unidentified species UI01, respectively. Low concentrations of As^V^ and two additional unidentified As species (UI03 and UI04) were also detected. Oligo-MM^12^ CecC had significantly different As species composition than germ-free CecC, most notably the absence of DMAs^V^ and UI04, along with the appearance of an unknown, oxidation sensitive species UI02. UI01 and UI03 were detected in all three mouse groups, but their mean concentrations were significantly higher in the SPF mice than in germ-free mice (UI01: adjusted p-value = 0.018; UI03: adjusted p-value = 0.004; Unpaired t test with Welch correction on log-transformed data). Notably, As thiolation in the form of dimethylmonothioarsinic acid (DMMTAs^V^) was seen exclusively in the CecC of Oligo-MM^12^ and SPF mice, and therefore microbiota-dependent. Moreover, As^III^ was detectable in Oligo-MM^12^ and SPF, but absent from germ-free caecal contents. The main differences in SPF CecC relative to Oligo-MM^12^ CecC were the significantly lower concentrations of species UI02 (Adjusted p-value = 0.019; Unpaired t test with Welch correction on log-transformed data) and appearance of an additional unidentified As species UI05. None of the unidentified As species UI01-UI05 found in CecC matched the unidentified species UI00 already present in the diet (**Fig. 1b**). The possibility of any of the detected unidentified species to correspond to trimethylarsine oxide (TMAs^V^O), MMAs^V^, monomethylarsenous acid (MMAs^III^), or dimethyldithioarsinic acid (DMDTAs^V^) could be dismissed based on either spiking of the samples with standards or by oxidation and re-analysis by both ion pairing and cation exchange chromatography (see **Supplementary material: Section II**).

These findings reveal a correlation between microbiota-dependent *in vivo* biotransformations of dietary AB and an elevated total As accumulation in the intestinal lumen. Notably, the dietary AB reached the large intestinal compartment, where the observed marked changes in As speciation suggest metabolization of AB through distinct host and microbial pathways. Substantial amounts of the As from AB-rich diet were transformed into DMAs^V^, a low-accumulating and lowly toxic form, in germ-free mice. In contrast, colonized mice exhibited higher levels of the more toxic and persistent arsenic species, As^V^ and As^III^. This observation prompted us to hypothesize that the systemic accumulation and persistence of arsenic from an AB-rich diet may consequently be influenced by gut microbiota status.

### Microbiota-dependent As transformation is associated with increased tissue accumulation of As from AB-rich diet

Figure 2 summarizes the total As concentrations in the blood circulation and various organ tissues of these mice. Correlating with the observed As accumulation in the intestinal compartments (see Fig. 1c), the As concentrations in lung, kidney, gall bladder, liver, and whole blood of conventional SPF mice were significantly increased in comparison to germ-free animals. Colonization of the simple 12-species bacterial consortium in the gnotobiotic Oligo-MM^12^ mice could not recapitulate this phenotype. In contrast to the other body tissues studied, As accumulation in the brain appeared to be independent of microbiota status. Notably, the most extreme As concentrations were exhibited in the gall bladders of SPF mice.

**Figure 2.**
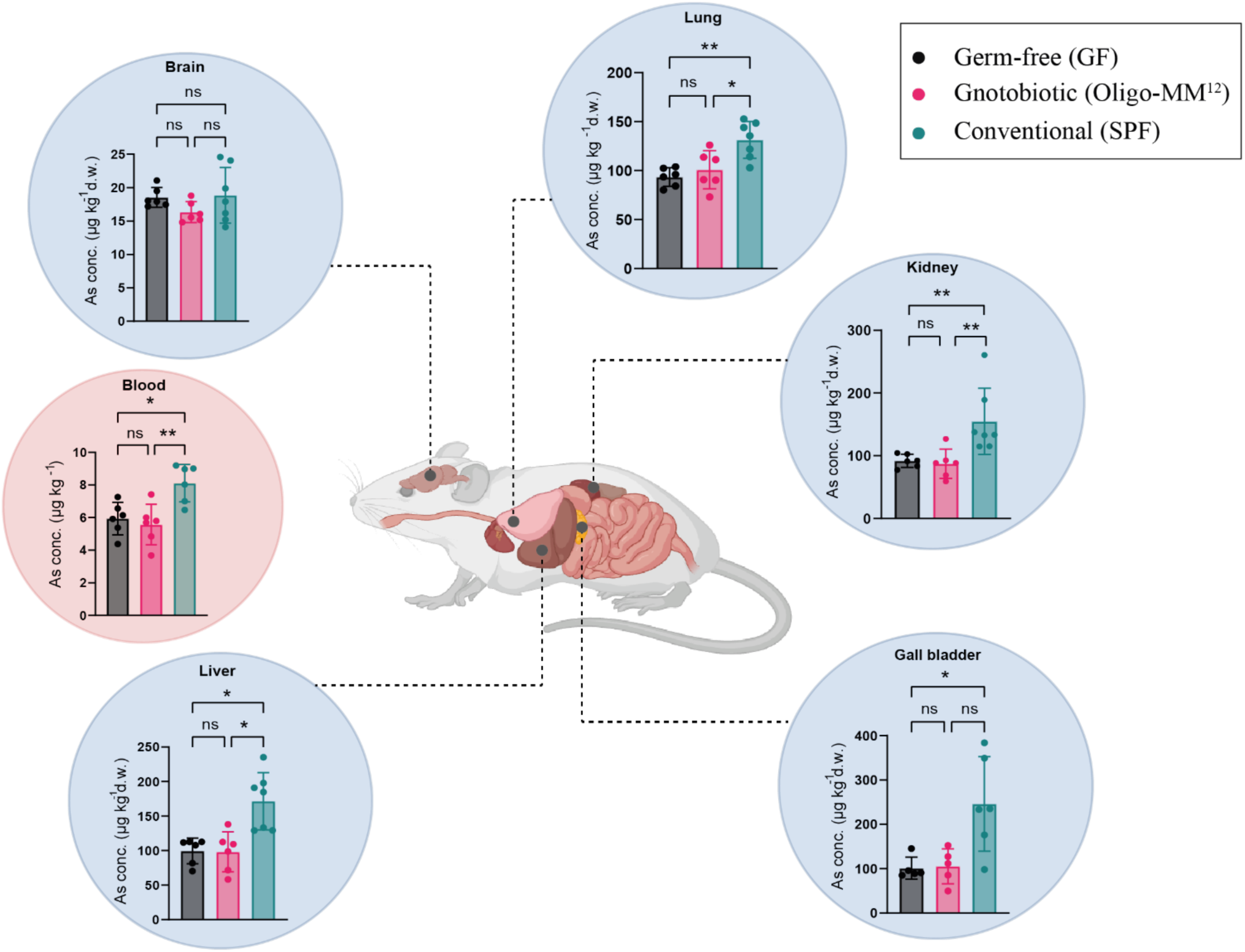
As distribution in the body compartments of mice of different microbiota statuses post chronic exposure. Samples were collected from germ-free (n=6), Oligo-MM^12^ (n=6), and SPF (n=7) mice after six weeks of feeding with the AB diet. Liver, gall bladder, kidneys, lungs, brain, and blood were sampled from all the mice. As concentrations are reported per kg of the dry weight of the samples, except for blood, which is reported per kg of the liquid blood. Groupwise mean values are represented in each of the graphs along with individual data points. The error bars represent the standard deviation from the mean value. The normality of each of the data sets was determined with the Shapiro-Wilk test. One-way ANOVA was performed to identify significant differences between the mean values of each of the mouse groups. (Lung: **, adjusted p-value = 0.0024; *, adjusted p-value = 0.0129; ns, adjusted p-value = 0.7186. Blood: **, adjusted p-value = 0.004; *, adjusted p-value = 0.0124; ns, adjusted p-value = 0.8402.) A non-parametric Kruskal-Wallis test was done to determine significant differences in the liver and gall bladder samples. (Liver: Germ-free vs. conventional, *, adjusted p-value = 0.0158; Gnotobiotic vs. conventional, *, adjusted p-value = 0.0134; ns, adjusted p-value > 0.9999. Gall bladder: *, adjusted p-value = 0.0298; Germ-free vs. gnotobiotic, ns, adjusted p-value > 0.9999; Gnotobiotic vs. conventional, ns, adjusted p-value = 0.0769). Ordinary one-way ANOVA was performed on the log-transformed data sets for the kidney (Germ-free vs. conventional, **, adjusted p-value = 0.0067; Gnotobiotic vs. conventional, **, adjusted p-value = 0.0022; ns, adjusted p-value = 0.8641). The SPF mice show significantly higher As concentrations compared to the germ-free mice in all the compartments except the brain. The As levels detected in the Oligo-MM^12^ mice are much more comparable to the germ-free mice in all the samples except for the brain where the levels are comparatively lower than the germ-free mice.

### SPF colonization status delays clearance of the accumulated As from the body

Our data indicated that complex microbiota colonization amplifies the accumulation of As in the intestines, faeces, and tissues during chronic consumption of an AB-rich diet. This observation led us to hypothesize that SPF mice with a complex microbiota would exhibit a slower As clearance kinetic compared to germ-free mice. To test this hypothesis, we transitioned a sub-cohort of mice from chronic As-rich diet consumption to a purified low-As diet containing only trace levels of As (16.2 ± 0.9 µg kg^−1^ total As; **Supplementary Fig. S1**). After a two-week clearance period, we measured total As concentrations in whole blood, livers, gall bladders, kidneys, lungs, brains, and intestinal contents of germ-free, Oligo-MM^12^, and SPF mice (Fig. 3). A two-week clearance period was decided to be sufficient based on data obtained from a pilot study monitoring faecal As excretion (**Supplementary Fig. S10**). At the end of the 14-day observation period, SPF mice exhibited elevated concentrations of total As remaining in intestinal contents, livers, gall bladders, lungs, and kidneys compared to germ-free and gnotobiotic mice. However, As levels in blood had equalized among all three mouse groups (compare Fig. 2 and Fig. 3), and like chronic accumulation (see Fig. 2), As retention in brain tissue was also microbiota status-independent.

**Figure 3.**
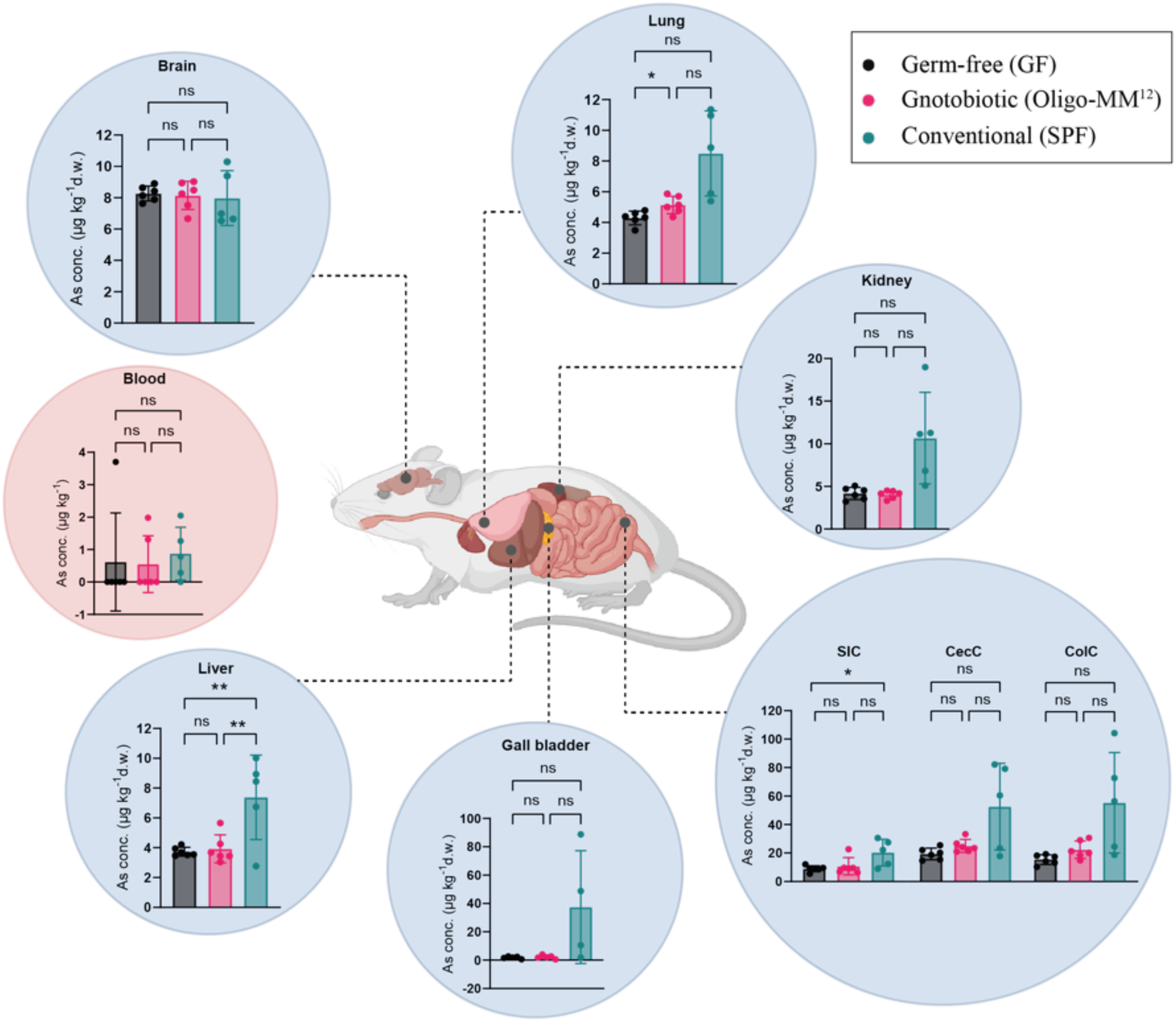
Body distribution of total As in mice of different microbiota statuses post a 2-week clearance period. Liver, gall bladder, kidney, lung, and brain tissues, and whole blood were sampled from germ-free (n=6), Oligo-MM^12^ (n=6), and SPF (n=7) mice following six weeks of feeding of AB-rich chow diet. As concentrations are reported per kg dry weight (d.w.), except for blood (per kg fresh weight). Bars represent groupwise mean values, and each scatter plot data point represents one mouse. Error bars indicate standard deviations. The normality of distribution was determined with the Shapiro-Wilk test. One-way ANOVA was performed to identify significant differences between the mean values of each of the mouse groups. (Lung: **, adjusted p-value = 0.0024; *, adjusted p-value = 0.0129; ns, adjusted p-value = 0.7186. Blood: **, adjusted p-value = 0.004; *, adjusted p-value = 0.0124; ns, adjusted p-value = 0.8402.) A non-parametric Kruskal-Wallis test was done to determine significant differences in the liver and gall bladder samples. (Liver: Germ-free vs. conventional, *, adjusted p-value = 0.0158; Gnotobiotic vs. conventional, *, adjusted p-value = 0.0134; ns, adjusted p-value > 0.9999. Gall bladder: *, adjusted p-value = 0.0298; Germ-free vs. gnotobiotic, ns, adjusted p-value > 0.9999; Gnotobiotic vs. conventional, ns, adjusted p-value = 0.0769). Ordinary one-way ANOVA was performed on the log-transformed data sets for the kidney (Germ-free vs. conventional, **, adjusted p-value = 0.0067; Gnotobiotic vs. conventional, **, adjusted p-value = 0.0022; ns, adjusted p-value = 0.8641).

Notably, the relative As concentration differences between SPF and germ-free/Oligo-MM^12^ mice observed during chronic exposure further increased during the As clearance phase (compare Figs. 2 and 3), indicating a delayed clearance of As in SPF mice.

This observation was further supported by monitoring the time courses of faecal As excretion from germ-free, gnotobiotic Oligo-MM^12^, and SPF mice for 14 days following the diet shift (Fig. 4). Initially, faecal As concentrations of the germ-free mice declined more slowly than in Oligo-MM^12^ and SPF animals. However, after the initial, rapid clearance phase, SPF mice exhibited a more intricate second clearance phase with a more complex kinetic. Starting on day 3, some SPF mice displayed intermittently re-increasing and prolonged faecal As excretion compared to the germ-free and Oligo-MM^12^ mice. The observed high interindividual variability of this delayed faecal clearance was consistent with the notable variability in intestinal and tissue retention of As among the SPF mice shown in Figure 3.

**Figure 4.**
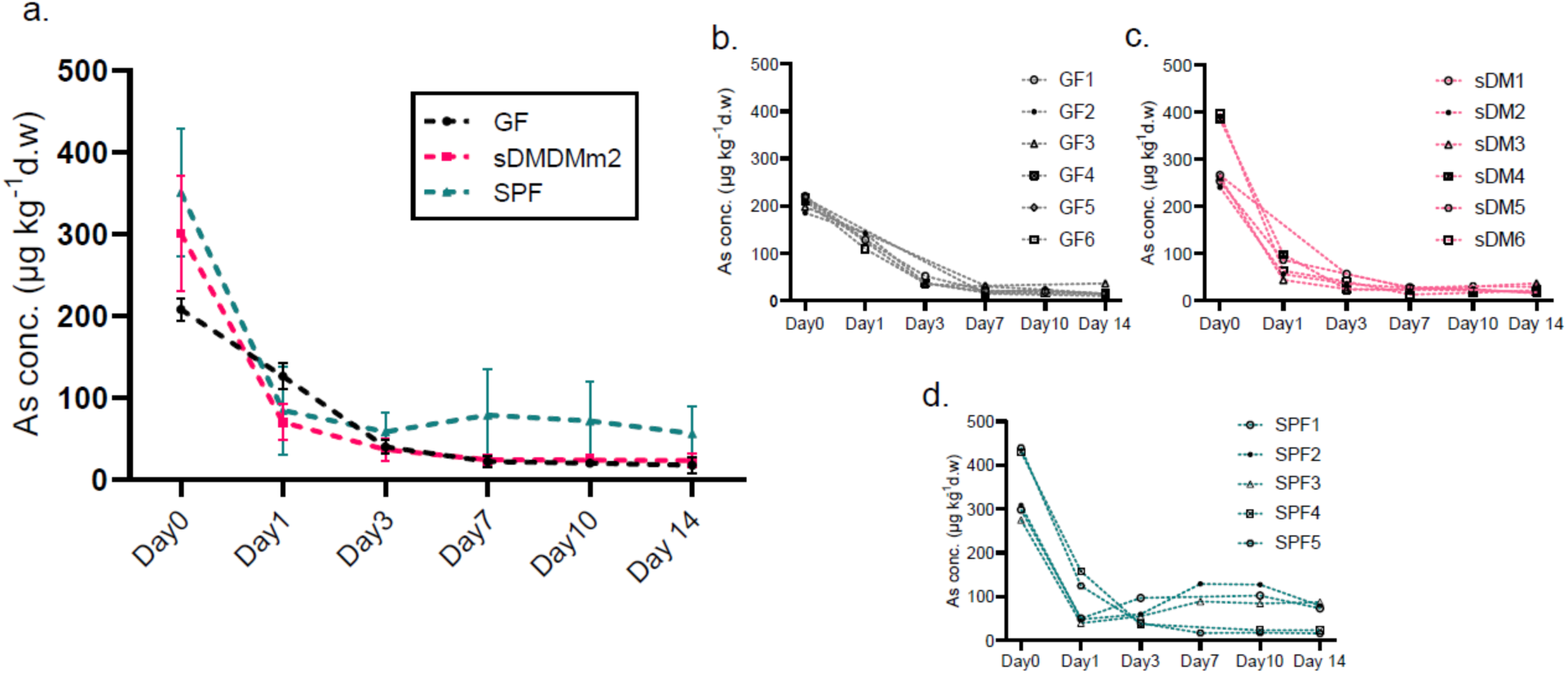
Faecal As excretion from germ-free, Oligo-MM^12^, and SPF mice during 14 days following diet shift from AB-rich diet to low-As-control diet. Faecal pellets were sampled from GF (n=6), Oligo-MM^12^ (n=6) and SPF (n=5) at days 0, 1, 3, 7, 10 and 14 post diet change. (a) Time course of mean±SD As concentrations. Symbols show means. Error bars indicate standard deviations. The number of replicates varied between the time points since all the mice produced faecal samples at every time point. (b-d) Time courses of faecal As concentrations in individual GF (panel b), Oligo-MM^12^ (panel c; “sDM”), and SPF (panel d) mice. The following data points could not be measured due to unavailability of faecal samples: on day 1, GF3, -4, -5 and sDM1; on day 3, GF3, -4, and sDM5; on day 7, SPF1 and -4; on day 10, GF3 and -4, sDM1 and -3.

## Discussion

Given its current non-toxic classification, there are no regulatory limits for AB in seafood. However, several studies have provided evidence for *in vivo* metabolism of AB [51, 55, 56], but did not definitively establish the causal roles of intestinal bacteria or host in the observed AB transformations. Our study addresses this gap by examining the murine *in vivo* transformation of As from a naturally AB-rich diet leveraging gnotobiology to distinguish host from microbial contributions to AB-metabolism.

Our data show that chronic As exposure through an AB-rich diet results in microbiota-dependent As accumulation within the intestinal lumen, likely attributed to microbial uptake of As, as reported for diverse environments such as water, soil, and the fish intestine [16, 17, 57, 58]. Additionally, we conducted a detailed characterization of the large intestinal As metabolite profile of mice subjected to chronic feeding of an AB-containing diet, revealing an unprecedented spectrum of As species depending on microbial colonization status.

Our study challenges the notion that AB remains unmetabolized *in vivo* as reported in prior studies in rodents or humans. Prior toxicologic studies often relied on single-dose exposures of AB, either in pure form in solution or within an AB-containing (seafood) meal [42–45, 47]. While an *in vitro* study reported the human gut microbial degradation of AB into DMAs^V^, DMAA (dimethylarsinoylacetate), and TMAs^V^O [30] after ≥7 days of incubation, such a prolonged timeframe exceeds the average human (and murine) gut passage time and therefore is unlikely relevant for single *in vivo* exposures. However, persistent dietary exposure entails recurrent dosing, extending the availability of AB to gut microbes and potentially fostering microbial transformation of AB.

Our work, analogous to a recent chronic feeding study in mice involving fish diet [51], reveals a diverse array of As species formed in mice chronically consuming AB-rich diet, including organic As species like DMAs^V^ and highly cytotoxic DMMTAs^V^ [59], as well as the toxic inorganic species As^V^ and As^III^, along with several unidentified species (see Fig. 1d). Two microbial pathways for AB degradation have been described to date, one leading to the formation of TMAs^V^O, and the other to DMAA. Both pathways eventually result in DMAs^V^, that may be further converted to As^V^ [60–62] – a common pathway of microbial As demethylation [30, 60, 63]. Notably, in the germ-free caecum, DMAs^V^ predominated, suggesting the possibility of host-mediated demethylation of AB into DMAs^V^ or a rather substantial accumulation of dietary DMAs^V^, that constituted merely 2% of the total As content in the AB-rich diet (see Fig. 1b).

In sharp contrast, in conventional SPF and gnotobiotic Oligo-MM^12^ mice the absence of DMAs^V^ and concomitant appearance of the thiolated As species DMMTAs^V^ strongly indicate a microbial role in their production. DMMTAs^V^ is a known *in vitro* murine microbial metabolite of DMAs^V^ [24], and As thiolation has previously been attributed to human and rodent gut microbial metabolism [18, 21, 64]. We propose that microbes in the SPF and Oligo-MM^12^ mouse gut produce DMMTAs^V^ from DMAs^V^ made available by host metabolism (and would otherwise accumulate in the germ-free mouse gut). Alternatively, these thiomethylated As species may be produced abiotically from DMAs^V^ in the presence of reduced sulfur compounds accumulated by anaerobic intestinal microbial metabolism [65].

Interestingly, elevated caecal concentrations of highly toxic As^III^ [66] were detected specifically in colonized mice, suggesting microbial demethylation of DMAs^V^ to As^V^ and subsequent microbial reduction [67, 68]. In the colonized, strictly anaerobic large intestinal lumen, highly reactive As^III^ is stabilized against spontaneous oxidation to As^V^, allowing for its local accumulation. Our data thus indicate that mammalian metabolism of As derived from the AB-rich diet predominately leads to DMAs^V^, which gut microbes might further thiolate or demethylate, producing additional As metabolites, some of which can contribute to an increased effective toxicity in the system [69].

Several As species detected in our samples remain unidentified. It is conceivable that DMAA could be among them, potentially serving as a precursor to DMAs^V^ according to the AB degradation pathway described above. Another potential candidate for an unidentified As species is TETRA (tetramethylarsonium ion), which might be generated through the decarboxylation of AB [70, 71]. Notably, TETRA has been detected in the urine of rats after oral AB consumption [55, 56]. Thus, TETRA emerges as a potential contributor to the pool of unidentified As species.

Our proposed *in vivo* transformations of As species are grounded in existing literature. It is essential to acknowledge the inherent complexity of *in vivo* As transformations, given the intricate interplay between host and microbial metabolic pathways. Nonetheless, our data unequivocally reveals distinct biotransformation pattern of dietary As in a microbially colonized gut compared to a germ-free gut. The array of As species detected, particularly DMMTAs^V^ and various unidentified species, suggests novel and unexplored AB metabolic pathways. Future experiments utilizing a purified diet containing AB as the sole As source could elucidate which As species are exclusively derived from dietary AB.

Our observation of significantly elevated As concentrations in livers, kidneys, lungs, gall bladders, and blood of conventional SPF mice (see Fig. 2) suggests higher systemic As exposure in these mice. This heightened exposure might result from microbiota-mediated increased bioaccessibility and bioavailability of ingested As [14, 17, 72, 73]. The increased As levels in SPF mice cannot solely be attributed to food intake differences, as germ-free conditions have been shown to prompt higher, rather than lower, food consumption to compensate for reduced calory extraction in absence of a gut microbiota [74, 75]. Moreover, high levels of As in the SPF gall bladders indicate enterohepatic circulation of As metabolites to contribute to As toxicokinetics *in vivo* in these mice [76, 77].

The gnotobiotic Oligo-MM^12^ mice, despite being densely microbially colonized, exhibited As concentrations more akin to germ-free mice. This underscores that body accumulation of As from AB-rich diet is more dependent on microbiota composition or diversity than density, indicating a role for specific microbial functions absent in the simplified 12-species Oligo-MM^12^ microbiome.

Quantifying total As concentrations in the faeces provided an overview of microbial influence on As excretion (see **Fig. S9**). Chronically exposed germ-free mice exhibited comparatively lower faecal As excretion, possibly due to an efficient urinary As excretion, particularly in the forms of AB and DMAs^V^ (see Fig. 1d), lowering the As burden in these mice. The necessary strict bioexclusion precautioned in these experiments allowed only the generation of a small number of spot urine samples (data not shown), which prohibited the reliable estimation of urinary excretion kinetics. Provided that innovative metabolic cage systems can be established that would enable continuous urine sampling and anaerobic freeze-preservation from germ-free animals, future studies may be able to unveil the excretion pathways maintaining lower As burden in germ-free mice.

Increased intestinal luminal accumulation and faecal As excretion in Oligo-MM^12^ and SPF mice may result from As immobilization by bacterial adsorption or uptake. Previous studies have noted bacterial cell mass-associated faecal As excretion, where antibiotic-induced microbiota depletion led to decreased faecal As concentrations [15, 16]. It is important to note that increased faecal As excretion in SPF mice does not necessarily imply reduced accumulation in the various systemic body compartments. Two previously described phases of As elimination in mice involve a rapid α phase elimination from the gut and a delayed β phase elimination releasing As from tissues and organs. Repeated dosing, comparable to chronic exposure, akin to our study, has been shown to slow down the elimination rate compared to a single dose [78]. Concomitantly increased faecal excretion, along with body accumulation in SPF mice, may be attributed to β phase elimination, implying a higher systemic As exposure. Higher systemic exposure in SPF mice is supported by elevated As concentrations in the lungs, kidneys, and livers after a two-week clearance period (see Fig. 3). Since intestinal re-absorption and enterohepatic circulation of As have been demonstrated in rodents previously [76, 77, 79], we hypothesize that gut microbes in SPF mice might delay As clearance partly by enhancing entero-hepatic circulation under chronic exposure. Monitoring faecal As excretion during the clearance experiment revealed a delayed intestinal-faecal As release in SPF mice, possibly due to β elimination (see Fig. 4a). However, substantial interindividual variability was noted in these mice (see Fig. 4d), which may be attributable to inherent microbiota composition variability [10, 12, 15].

In conclusion, microbial metabolism of seafood-derived dietary As, at levels commonly found in food [80], potentially leads to the formation of toxic As species *in vivo*. This information is highly relevant for risk assessment of As from seafood and can be directly utilized by national and international regulatory authorities such as the World Health Organization or the European Food Safety Authority to ensure food safety. The toxicokinetics of the ingested As varied depending on the gut microbial complexity, and a diverse gut microbiota could delay As clearance during long-term consumption of a diet with environmentally relevant levels of AB [81]. Our study emphasizes the importance of considering the role of gut microbial metabolism when assessing chronic exposure to As through diet, particularly for lowly toxic As species such as AB.

## Materials and Methods

### Mouse maintenance

Animal experiments were performed under the animal experimental licenses BE44/18 and BE45/21 approved by the Bernese Cantonal Committee for Animal Experiments and carried out in accordance with the Swiss Federal Law for animal experimentation.

All mice used in this study were in the same C57BL/6J strain background. Germ-free mice were generated and maintained in flexible film axenic isolators at the Clean Mouse Facility (CMF) of the University of Bern (Switzerland). The germ-free status of all animals has been routinely monitored using culture-based and culture-independent methods established and validated by the CMF. Gnotobiotic Oligo-MM^12^-associated mice have been established at the CMF by inoculation of germ-free C57BL/6J mice with pure cultures of 12 fully genome sequenced murine intestinal bacterial species known as the Oligo-MM^12^ community [53] and stably maintained in flexible film isolators under strict axenic conditions [82]. For this study, isogenic gnotobiotic Oligo-MM^12^ mice were generated by co-housing of germ-free mice at 5-6 weeks of age with Oligo-MM^12^ colonizer animals for a span of 4 weeks. All germ-free and Oligo-MM^12^ mice used in this study were housed under axenic barrier conditions at the Clean Mouse Facility of the University of Bern.

Conventional specific pathogen-free (SPF) mice were maintained at the Central Animal Facility (CAF) of the University of Bern. To avoid between-facility mouse genetic variability, an isogenic founder population of SPF mice was derived from germ-free mice transferred from the CMF through co-housing with SPF colonizer mice maintained at the CAF for four weeks before expansion of the colony under SPF conditions. The experimental SPF mice were housed in autoclaved Sealsafe-plus IVC cages (Tecniplast, Italy).

At CMF and CAF, mouse rooms had controlled ambient temperature (23–25°C), relative humidity (50– 60%), and day/night cycle (12/12). Animals had access to food and drinking water *ad libitum*.

All gnotobiotic mice housed at the CMF (germ-free and Oligo-MM^12^ in this case) were routinely maintained on an autoclavable rodent chow diet Kliba-Nafag 3307 (Granovit, CH), while the SPF mice at the CAF were routinely maintained on irradiated diet Kliba 3432 (Granovit, CH). These diets were analysed for their total As concentration and Kliba-Nafag 3307 contained a total As concentration of 431 ± 14 µg kg^−1^ while Kliba 3432 contained 54 ± 7 µg kg^−1^ of total As.

Drinking water was provided *ad libitum* throughout the experiments. Drinking water was analysed and found to contain very low As concentrations of 5.18 ± 0.85 µg kg^−1^ and hence not expected to contribute to As exposure in our experiments.

### Chronic arsenic exposure

At the start of the experiment, age-matched (5-11 weeks old) female germ-free (n=12), Oligo-MM^12^ (n=12), and SPF (n=13) mice were shifted to a specific batch of the autoclaved (autoclaved at the same time in the same autoclave) rodent chow diet Kliba 3307 and fed with this diet *ad libitum* for six weeks. The product batch prepared for our study was analysed for total As concentration and As speciation using inductively coupled plasma-mass spectrometry and high performance liquid chromatography-inductively coupled plasma-mass spectrometry, respectively (for details see sections ‘***Total arsenic analysis***’ and ‘***Arsenic speciation analysis***’), and was found to be naturally contaminated with As at a total concentration of 431 ± 14 µg kg^−1^, of which 92 ± 2 % accounted for AB, 4.0 ± 0.2 % for As^V^, 2.1 ± 0.3 % for DMAs^V^, and the remaining 2.1 ± 0.1 % for an unidentified As species (UI00) (see Fig. 1b).

A subset (Germ-free, n=6; Oligo-MM^12^, n=6, and SPF, n=7) of mice were euthanized at the end of six weeks of feeding, and samples including liver, kidneys, lungs, brain, gall bladder and gut luminal contents like small intestinal, caecal and colonic contents were collected for As quantification. Retro-orbital blood and faeces samples were collected at a single time point shortly before euthanization. The remaining subset of mice (Germ-free, n=6; Oligo-MM^12^, n=6; and SPF, n=5) was kept alive to study As clearance kinetics following As chronic exposure.

### Arsenic clearance experiment

After chronic exposure to the As-rich diet, mice were switched to irradiation sterilized (2 X 20 kGy gamma followed by 30-60 kGy X-ray irradiation) open label purified diet D11112201i (Research Diets Inc., USA) that served as low-As control diet containing background As concentrations of 16.2 ± 0.9 µg kg^−1^ provided *ad libitum* for two weeks to allow for clearance of the previously accumulated As.

### Sample collection

Fresh faecal samples were collected and subsequently flash frozen in liquid nitrogen. Terminal retro-orbital blood sampling was done from each mouse to quantify the As concentration in the blood circulation. Organs, tissues, and intestinal contents were harvested post euthanization under sterile conditions and under an anaerobic atmosphere (Whitley Anaerobic Workstation A45, Don Whitley Scientific, UK) to preserve oxygen-sensitive As species. All the samples were collected in pre-weighed tubes and flash-frozen in liquid nitrogen shortly after collection and stored at −80°C until analysis. Five to six food pellets of the diet were sampled for As analysis from a larger batch that was later distributed among the experimental mice and the samples were stored at 4°C until analysis.

### Total arsenic analysis

Samples were processed and analysed for total As concentration at the Environmental Chemistry Laboratory (cLAB) from the Institute of Geography, University of Bern. The solid samples were freeze-dried (LyoQuest, Azbil Telstar) and ground into a powder. Diet samples were milled directly into a fine powder (RETSCH MM 400, Germany) and equally divided into subsamples. Pulverized solid samples and 100 µL of each of the liquid samples were weighed and microwave-digested with 0.3 ml of in-house doubly distilled, trace-metal grade 70% (w/w) nitric acid (HNO_3_, Grogg Chemie AG, Stettlen, Switzerland) and 0.2 ml of 30% (w/w) hydrogen peroxide (H_2_O_2_, Sigma-Aldrich). The microwave assisted acid digestion (MARS 6, CEM) was done according to the heating program below:

-A ramp time of 5 minutes to reach 35°C and a hold of 5 minutes at 35°C.

-A ramp time of 5 minutes to reach 55°C and a hold of 5 minutes at 55°C.

-A ramp time of 5 minutes to reach 75°C and a hold of 10 minutes at 75°C.

-A ramp time of 5 minutes to reach 95°C and a hold of 30 minutes at 95°C.

The digested samples were diluted with ultrapure water (18.2 MΩ cm) to obtain a final HNO_3_ concentration of 5% (v/v). Samples were then centrifuged (Multifuge X1, Thermo Fischer) at 1500 ×g for 10 min to separate the supernatant from the solid pellet. 100 µL of HPLC grade methanol (99.9%) was added to 2 mL of each of the sample supernatants before analysis by inductively coupled plasma-mass spectroscopy (ICP-MS, 7700x, Agilent Technologies) using an ASX-500 autosampler (Agilent Technologies). Calibration standards ranging up to 100 µg kg^−1^ and calibration checks at the low, mid, and high points of the calibration curve were prepared by weight from a multielement stock solution (IV-ICPMS-71A; Inorganic Ventures) in 5% (v/v) HNO_3_ and 5% (v/v) methanol. The certified reference materials (CRM) – BCR 185R bovine liver (IRMM, JRC, Belgium), ERM-BB184 bovine muscle (IRMM, JRC, Belgium), SRM 955c caprine blood (NIST, Japan) were also processed and analysed together with samples to determine the As extraction efficiency. The extraction efficiency of As from the CRM samples for the ‘Chronic arsenic exposure’ data were as follows – BCR 185R bovine liver: 77.50 ± 13.78 % and SRM 955c caprine blood: 100.29 ± 1.48 %. The extraction efficiency of As from the CRM samples for the ‘Arsenic clearance experiment’ data were as follows – BCR 185R bovine liver: 87.16 ± 5.98 %, ERM-BB184 bovine muscle: 77.53 ± 3.42 %, and SRM 955c caprine blood: 91.20 ± 3.96 %. Method blanks were also included to account for potential As contamination during sample processing and analysis.

### Arsenic speciation analysis

As species from the chow diet and murine caecal content samples were extracted under anaerobic conditions in a glovebox (Systemtechnik, Germany) with a deoxygenated mobile phase consisting of 5 mM tetrabutylammonium hydroxide (TBAH; Sigma-Aldrich), 3 mM malonic acid (99%, Alfa Aesar) and 5% (v/v) methanol (pH adjusted to 5.9). These samples were freeze-dried (LyoQuest, Azbil Telstar) and ground into a powder before the extraction. As speciation analysis was done by high performance liquid chromatography-inductively coupled plasma-mass spectrometry (HPLC-ICP-MS) with a 1260 Infinity HPLC coupled to a 7700 ICP-MS (both Agilent Technologies). Besides As (m/z 75), vanadium (m/z 51) was also monitored. Chloride ions form adducts with argon (^40^Ar^35^Cl) that can interfere with the signal for ^75^As. Chloride also forms an adduct with oxygen (^35^Cl^16^O) with the same m/z as ^51^V. Vanadium was thus monitored to discard any peak eluting as ^75^As that may have come from the presence of chloride-based salts in samples. As species were separated by ion-pairing chromatography using a Zorbax SB-C18 column (150 mm × 4.6 mm, 5 μm particle size, Agilent) through isocratic elution for 25 min with the mobile phase used for extraction. The column was maintained at 50°C and the flow rate was set as 1.2 mL min^−1^. This method was adapted from a previous study [83]. Calibration standards using DMAs^V^ were prepared in the mobile phase at As concentrations ranging from 0.1 to 50 μg kg^−1^. BCR 185R bovine liver CRM and method blanks were also processed and analysed with the samples. The extraction efficiency of As from the BCR 185R bovine liver CRM was 48 ± 3 % and the column recovery was 29 ± 3 %. Commercially available standards of arsenate (As^V^) (Sigma-Aldrich), arsenous acid (As^III^) (Sigma-Aldrich), monomethylarsonic acid (MMAs^V^) (Sigma-Aldrich), dimethylarsinic acid (DMAs^V^) (Sigma-Aldrich), trimethylarsine oxide (TMAs^V^O) (Toronto Research Chemicals, Ontario, Canada), arsenocholine (AC) (Toronto Research Chemicals, Ontario, Canada), and arsenobetaine (AB) (Toronto Research Chemicals, Ontario, Canada) were used for As species identification and as mix for chromatography optimisation. Standards of dimethylmonothioarsinic acid (DMMTAs^V^), dimethyldithioarsinic acid (DMDTAs^V^), and monomethylarsonous acid (MMAs^III^) were synthesized in-house at the cLAB, following previously established methods [84, 85]. To confirm the identity of co-eluting As species, samples were spiked with certified standards or oxidized with H_2_O_2_, or analysed through cation exchange chromatography using a PRP-X200 column (150 × 4.6 mm, 10 μm particle size; Hamilton, US) and 2.5mM pyridine (99.8%, Sigma-Aldrich) as mobile phase and the pH adjusted to 2.5-2.7 with the help of formic acid, following a method adapted from the literature [86].

### Chromatogram peak integration

Integration of chromatogram peaks to determine area under the curve and subsequently, compound concentrations was done using MassHunter 4.3 Workstation Software, 7700 ICP-MS Data Analysis (Version C.01.03).

### Statistical analyses

All statistical analyses were performed using GraphPad Prism version 9.3.1 for Windows (GraphPad Software, San Diego, California USA, www.graphpad.com). The normal distribution of data sets was checked with the Shapiro-Wilk test. Significant differences between the means of normally distributed data sets were determined with the help of ordinary one-way ANOVA. If datasets had significantly different variances, Brown-Forsythe and Welch ANOVA tests were used to determine significant differences between the means. In case of data sets not following a normal distribution, the non-parametric Kruskal-Wallis test was used to determine statistically significant differences between group means.

### Figures

Data graphs were generated using GraphPad Prism version 9.3.1 for Windows (GraphPad Software, San Diego, California USA, www.graphpad.com). Figures were prepared with Adobe Illustrator CS6. Figures 2 and 4 were illustrated utilizing BioRender.com.

## Acknowledgments

We thank Miquel Coll-Crespi and Dr. Amrika Deonarine for standardizing and developing an effective method for arsenic detection and quantification that was used for generating the data in this manuscript, and Rizlan Bernier-Latmani (EPFL, Switzerland) for the initial idea leading to this study. We thank the staff and management team of the Clean Mouse Facility and the Central Animal Facility of the University of Bern and the Genaxen foundation for animal experimental support. We thank all lab members of the Soil Science Group at the Institute of Geography and the Hapfelmeier lab at the Institute for Infectious Diseases at University of Bern for their support and advice. This project was funded by the University of Bern through the Interfaculty Research Cooperation “One Health” (https://www.onehealth.unibe.ch/).

## Supplementary material

### SECTION I

#### Experimental outline

**Figure S1.**
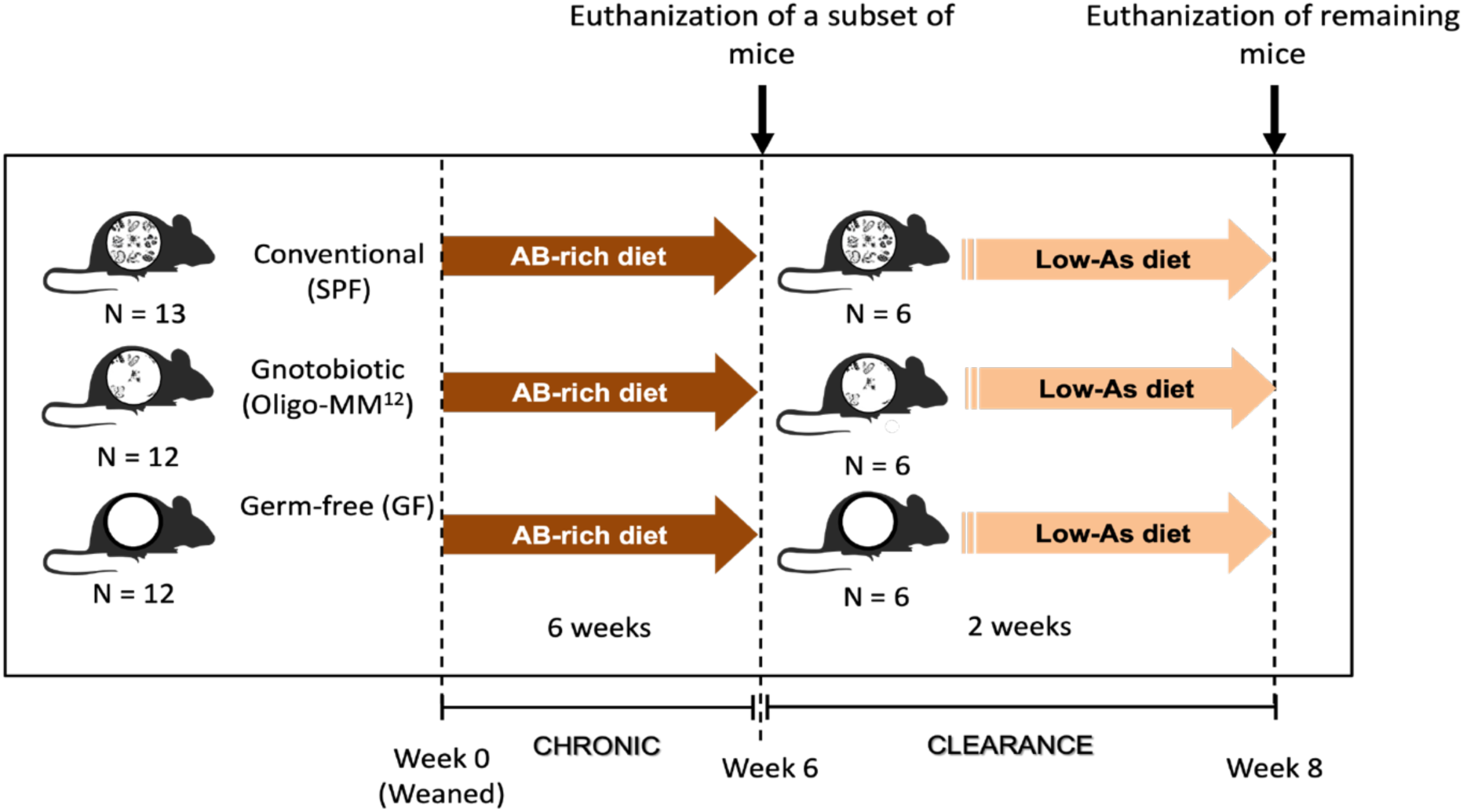
Experimental design. Age-matched (5-11 weeks old) female SPF (n=13), Oligo-MM12 (n=12), and germ-free (n=12) mice were fed on the same batch of autoclave sterilized rodent chow diet Kliba-Nafag 3307 that naturally contained As (‘AB-rich diet’) for six weeks. A subset (SPF, n=7, Oligo-MM12, n=6, and Germ-free, n=6) of the total number of mice were euthanized at the end of six weeks that constituted a period of chronic dietary As exposure. The remaining subset (SPF, n=6, Oligo-MM12, n=6, and Germ-free, n=6) of mice were switched to a low-As purified control diet and fed with this diet for two more weeks before euthanization to study the clearance kinetics of As.

### SECTION II

#### Identification of chromatogram peaks

This section is associated with Fig.1b and further elaborates on the identification/characterization of the As species. Both ion-pairing and cation exchange chromatography techniques were used for this purpose.

1. Ion-pairing chromatography:

A. **Unidentified As species UI01**: Assessing the Retention time (Rt) and based on available literature [87, 88], that states arsenocholine (AC) as a precursor of AB, AC seemed to be a potential candidate. Therefore, standard of AC was run along with the experimental samples on the HPLC using ion-pairing chromatography. Although AC co-eluted with the UI01 peak (**Suppl. Fig. S2**), UI01 could not be confirmed as AC because of the possibility of co-elution of all the unretained As species within this peak due to its early elution [Retention time (Rt) ∼1.915].
B. **Unidentified As species UI02 and UI03**: UI02 and UI03 likely corresponded to reduced, unstable species or thiol-containing As species based upon oxidation experiments (**Suppl. Figures S4-S6**).
C. **Unidentified As species UI04 and UI05**: UI04 in germ-free mice was found to be oxygen-stable (**Suppl. Fig. S4**), while species UI05 found in SPF mice may have been converted by oxidation to UI04 (**Suppl. Fig. S6**).
D. **DMMTAs^V^**: The presence of DMMTAs^V^ in the Oligo-MM^12^ and SPF CecC samples was confirmed by comparing the retention times (Rt) of the chromatogram peaks with that of a DMMTAs^V^ standard (**Suppl. Fig. S3**) and in oxidation experiments where DMAs^V^ (known oxidation product of DMMTAs^V^) appeared in the analysed samples following oxidation (**Suppl. Fig. S4-6**).
2. Cation exchange chromatography:

A. **AB**: All the mouse samples seemed to have AB (**Suppl. Fig. S8**) when compared to the As species standards (**Suppl. Fig. S7**).
B. **AC**: Small distinguishable peaks were seen at a position comparable to the AC standard (**Suppl. Fig. S7**) in the Oligo-MM^12^ and SPF sample chromatograms (**Suppl. Fig. S8**). However, no such peaks, distinguishable from the background noise could be detected in the germ-free sample chromatogram (**Suppl. Fig. S8**).
C. **As^III^**: The presence of As^III^ was confirmed in Oligo-MM^12^ and SPF mice but not in germ-free mice (**Suppl. Fig. S8)** when compared to the As standards (**Suppl. Fig. S7**).
D. **DMAs^V^, As^V^ and DMMTAs^V^**: These three As species were very difficult to distinguish from each other due to their similar peak positions on the chromatogram (**Suppl. Fig. S7**). Therefore, it was difficult to identify these three As species in the GF, Oligo-MM^12^ and SPF samples, all of which seemed to have a peak at that position on their respective chromatograms that might correspond to either one or a combination of these As species (**Suppl. Fig. S8**).
E. **Unidentified species**: Running samples in the cation exchange chromatography showed one unidentified species, UIC02, in germ-free and SPF CecC and another species, UIC01, in Oligo-MM^12^ mouse CecC (**Suppl. Fig. S8**) that did not correspond to any of the As species standards (**Suppl. Fig. S7**).

**Figure S2.**
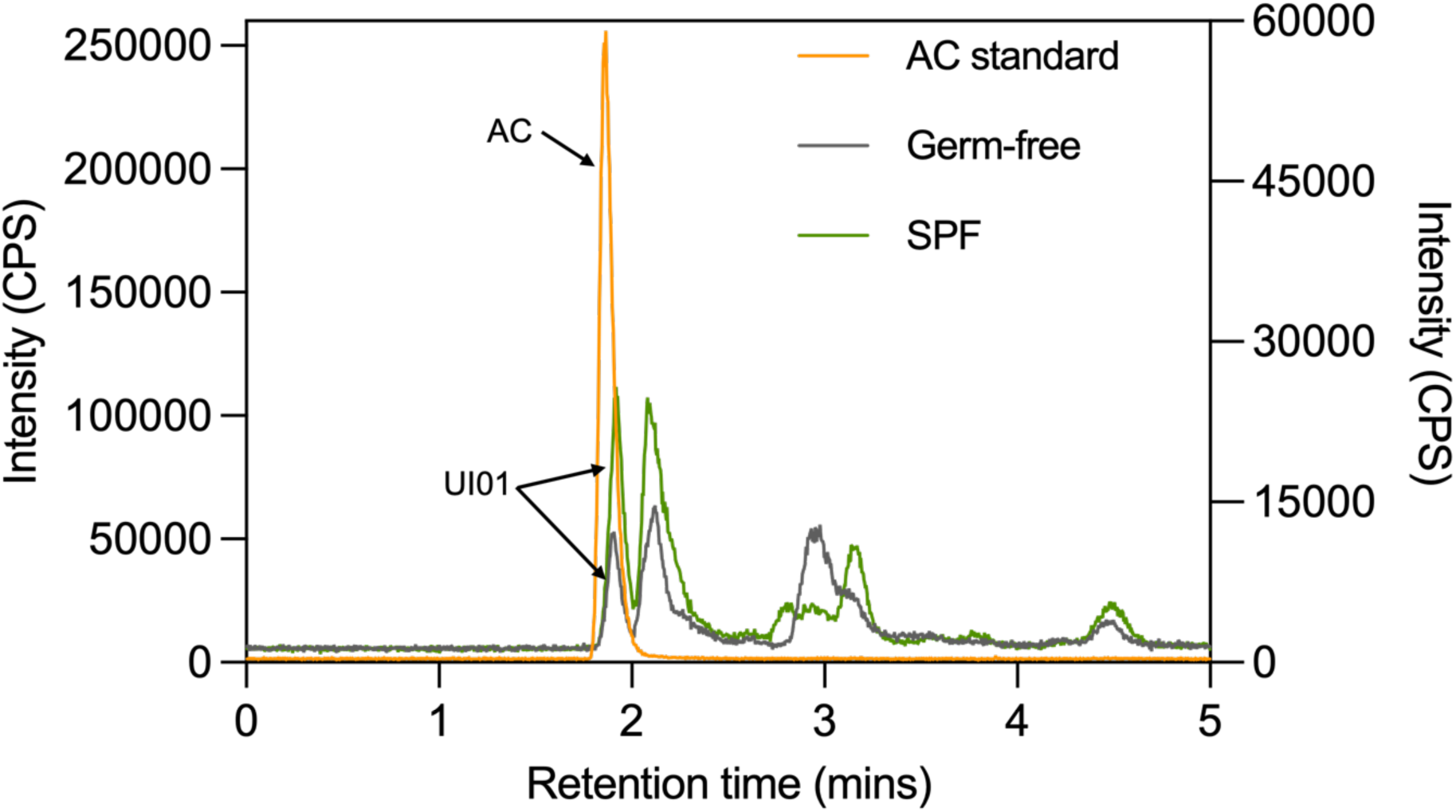
Ion pairing chromatograms of germ-free (GF) and conventional (SPF) caecal contents compared to the chromatogram of the standard of arsenocholine (AC). The left Y-axis represents the scale for Intensity (CPS: counts per second) for the AC standard chromatogram while the right Y-axis represents that for the GF and SPF sample chromatograms. The X-axis represents the retention time in minutes.

**Figure S3.**
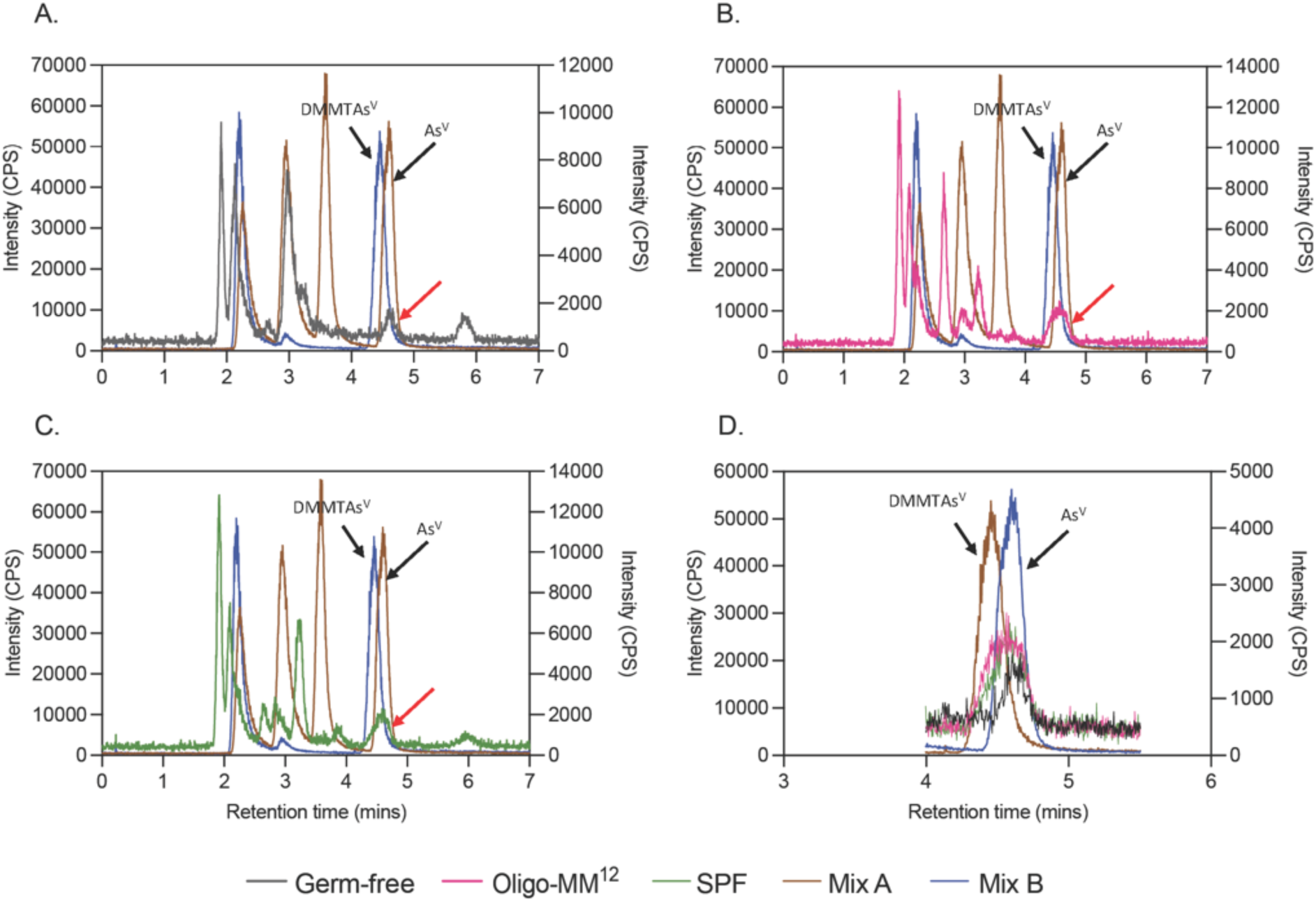
Ion pairing chromatograms obtained from germ-free (GF) [panel A], Oligo-MM^12^ [panel B], and conventional (SPF) [panel C] caecal content samples are compared to that of As standards in MixA (a mix of the standards of TMAsO, DMAs^V^, MMAs^V^, and As^V^) and Mix B (a mix of the standards of As^III^, DMMTAs^V^ and DMDTAs^V^). Black arrows indicate the peaks of DMMTAs^V^ and As^V^ in the standard mixes. The left Y-axis in each of the graphs (A-D) represents the scale for Intensity (CPS: counts per second) for Mix A and B chromatograms while the right Y-axis represents that for the sample chromatograms and the X-axes for all the graphs represent the retention time in minutes. The Oligo-MM^12^ [panel B] and conventional (SPF) [panel C] sample peaks (indicated by red arrow) seem to be a combination of both DMMTAs^V^ and As^V^ peaks while, the germ-free [panel A] peak (indicated by red arrow) align more to the As^V^ standard peak. Panel D shows the peak of interest from the three sample types (GF, Oligo-MM^12^ and SPF) overlaid by the DMMTAs^V^ and As^V^ peaks in the standard mixes.

**Figure S4.**
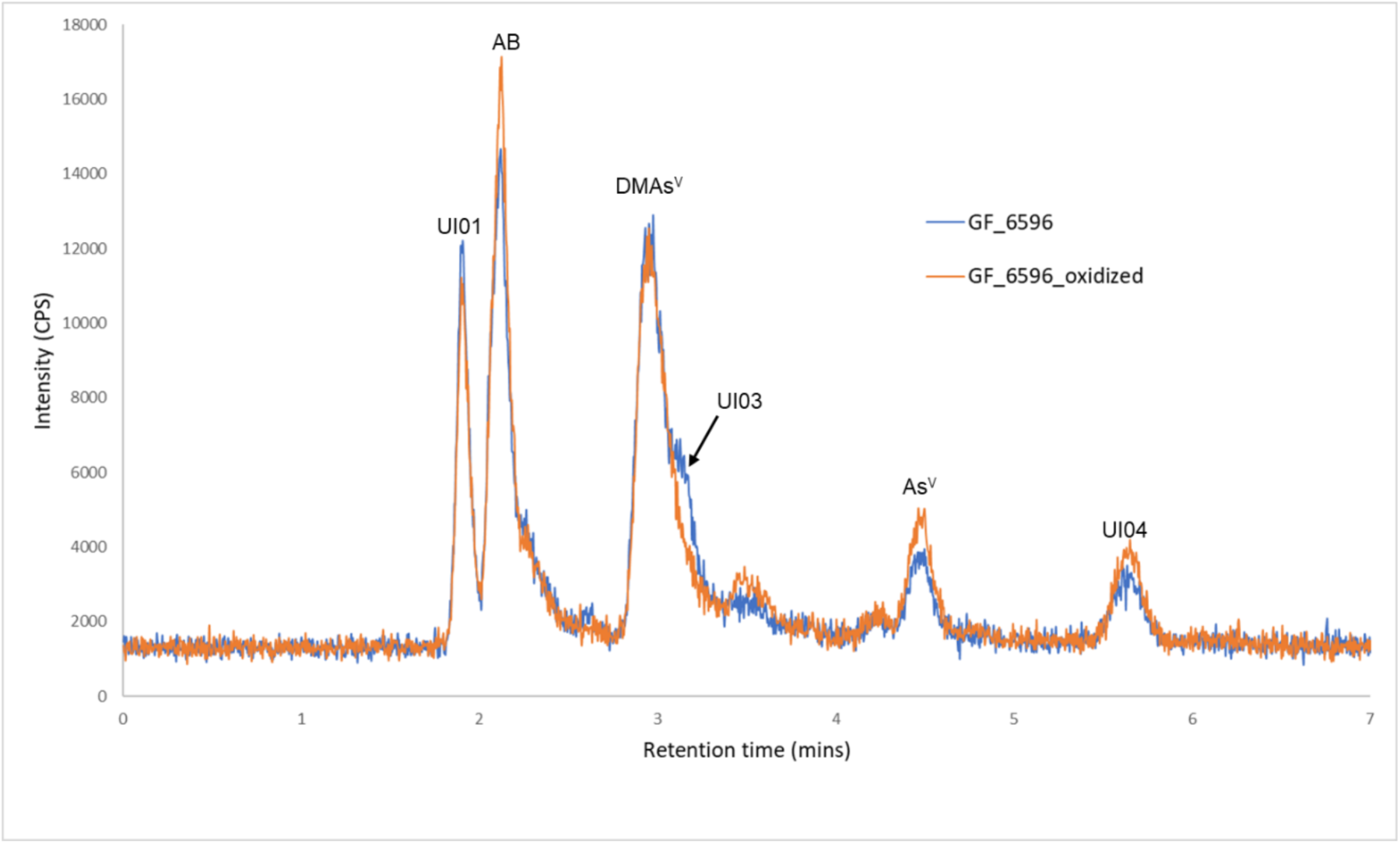
Ion pairing chromatograms obtained from the germ-free caecal content (blue) and its subsample subjected to oxidation (orange). The Y-axis represents the scale for Intensity (CPS: counts per second) and the X-axis represents the retention time in minutes. Three unidentified species namely, UI01, UI03 and UI04 were identified in the germ-free sample, among which UI03 seemed to decrease on oxidation. CPS: counts per second.

**Figure S5.**
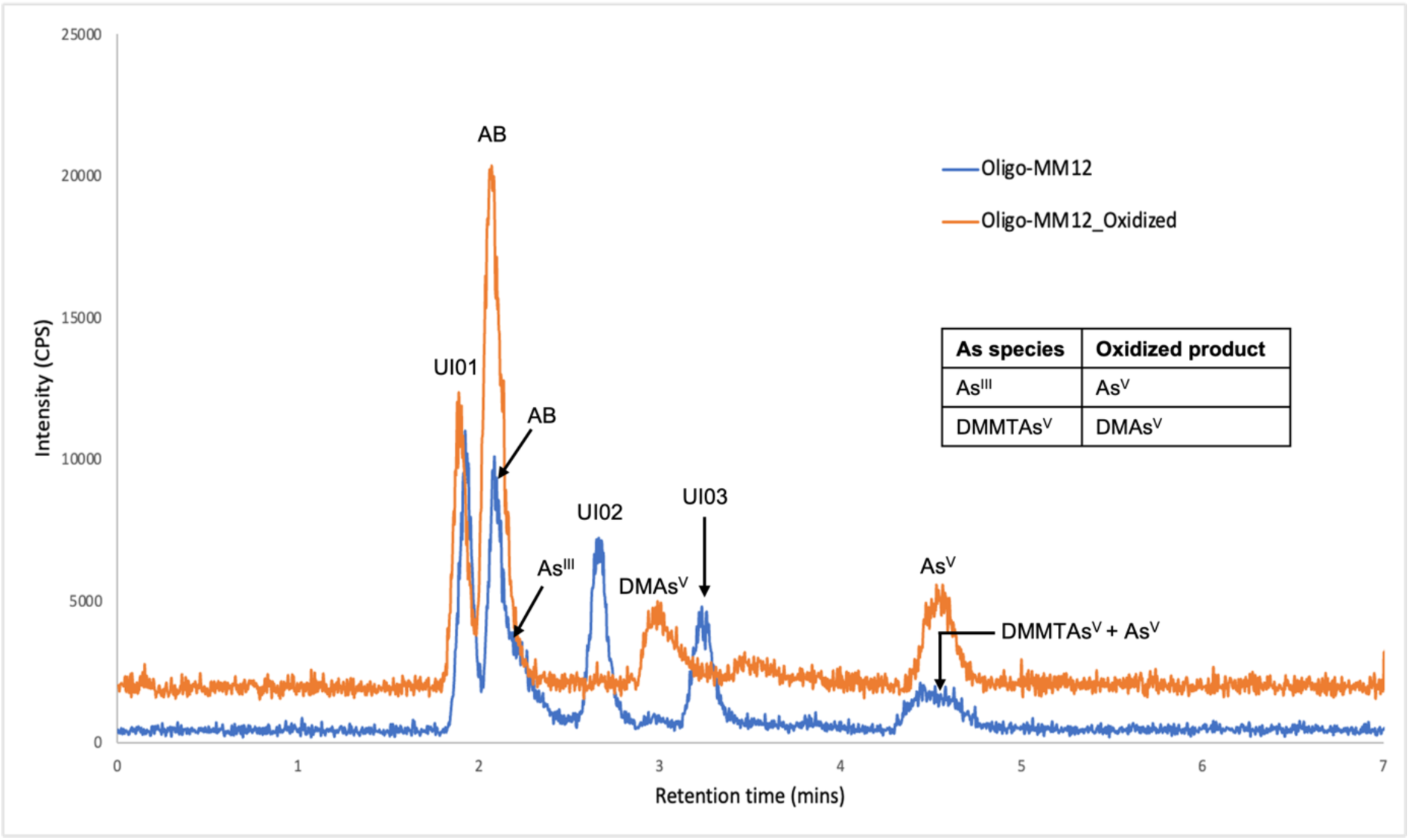
Ion pairing chromatograms obtained from the Oligo-MM^12^ caecal content (blue) and its subsample subjected to oxidation (orange). The Y-axis represents the scale for Intensity (CPS: counts per second) and the X-axis represents the retention time in minutes. In contrast to the germ-free mice, DMAs^V^ and UI04 are not seen in the Oligo-MM^12^ sample. However, a new unknown peak UI02 is detected that disappears on oxidation. Although As^V^ and DMMTAs^V^ co-elute forming a broad peak, the increase in DMAs^V^ levels upon oxidation indicates the presence of DMMTAs^V^ in the original sample. The table shows the expected oxidized products of some As species.

**Figure S6.**
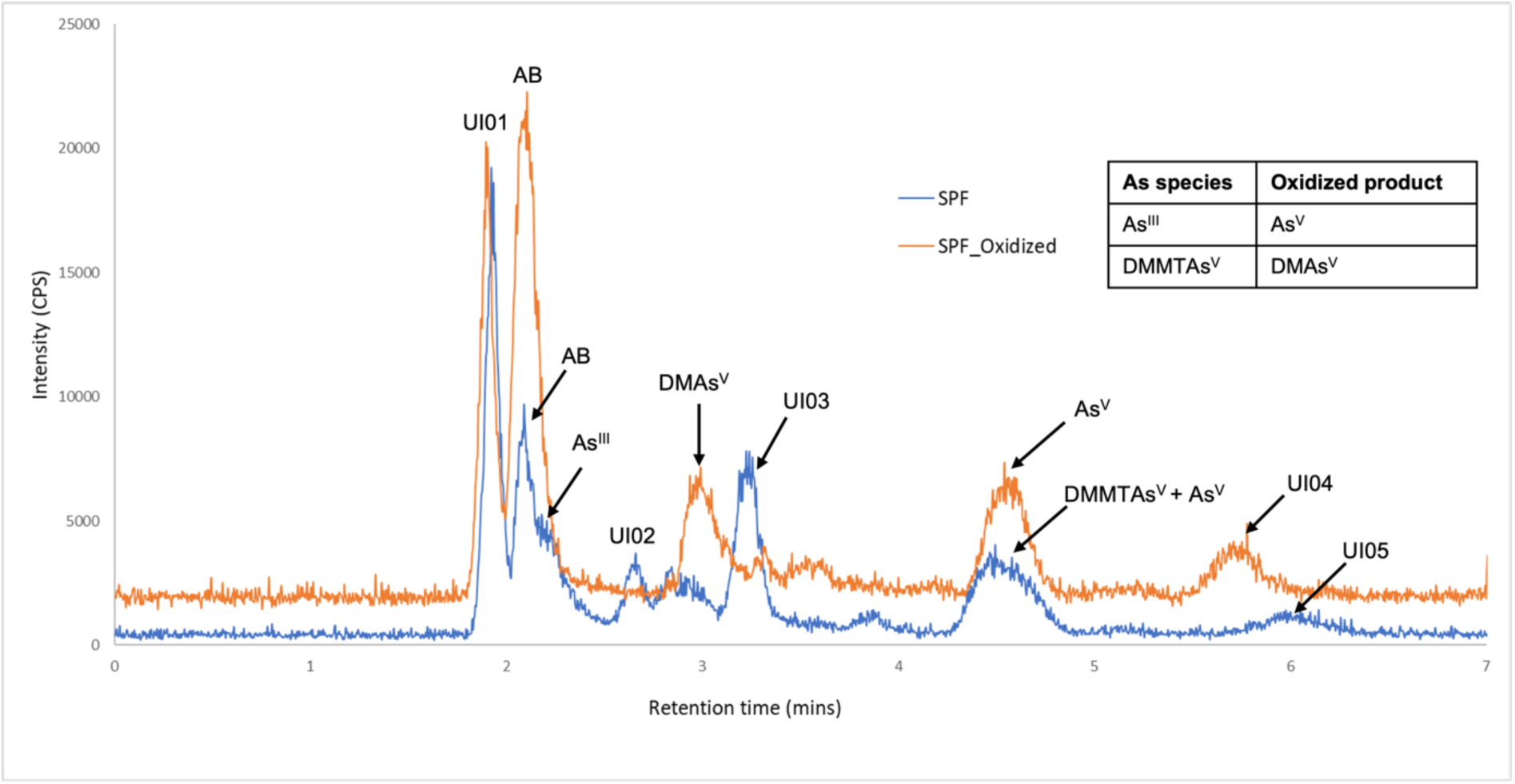
Ion pairing chromatograms obtained from the conventional (SPF) caecal content (blue) and its subsample subjected to oxidation (orange). The Y-axis represents the scale for Intensity (CPS: counts per second) and the X-axis represents the retention time in minutes. A completely new unidentified species UI05 was detected in the SPF caecal content. UI04 seen in the germ-free and not in the Oligo-MM^12^ caecal content was detected in the oxidized sample. Similar to the Oligo-MM^12^ samples, the presence of DMMTAs^V^ was indicated by the increased DMAs^V^ levels on oxidation. The table shows the expected oxidized products of some As species. CPS: counts per second.

**Fig. S7.**
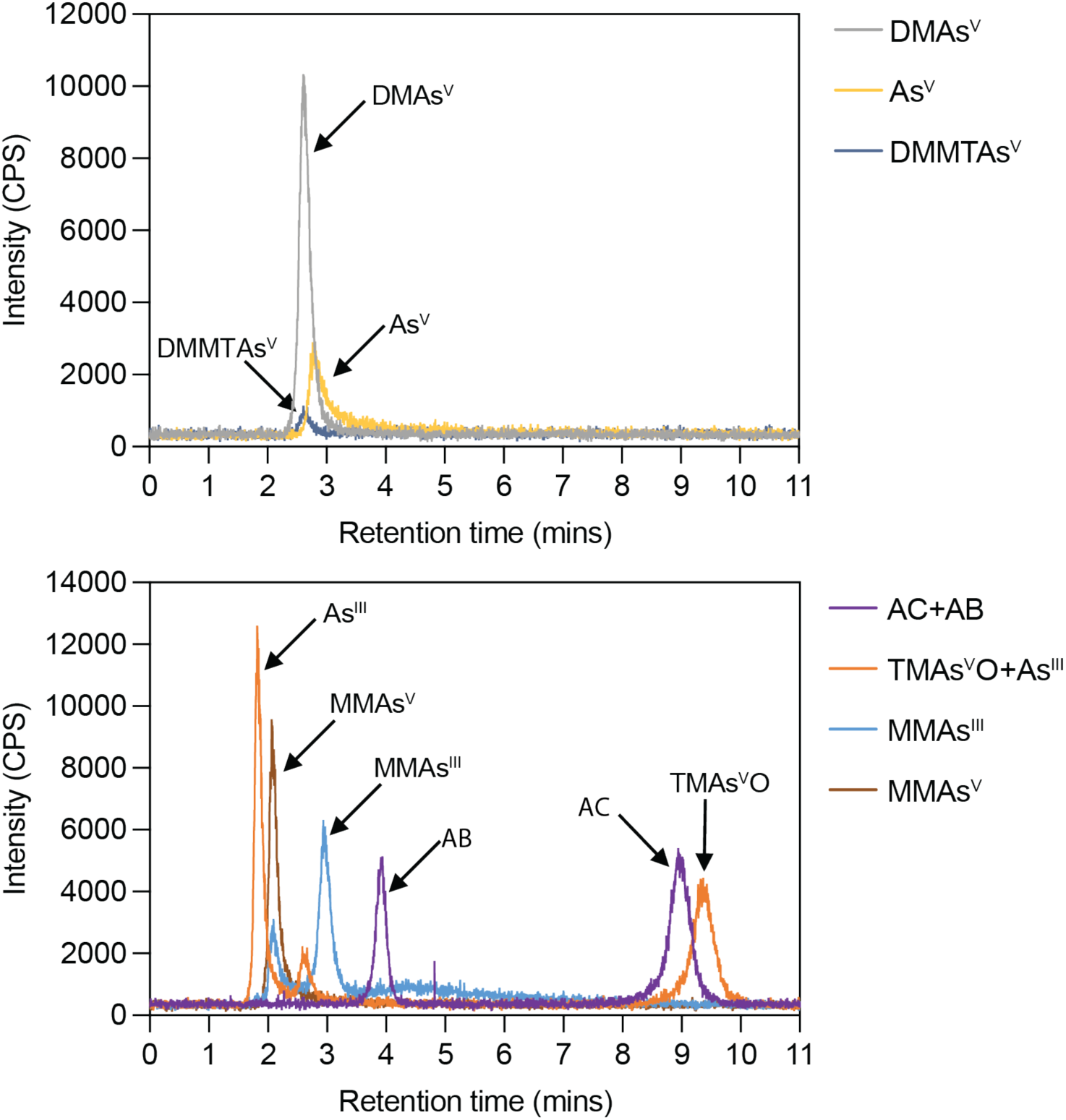
Cation exchange (CATEX) chromatogram of the As species standards. The Y-axis represents the scale for Intensity (CPS: counts per second) and the X-axis represents the retention time in minutes.

**Fig. S8.**
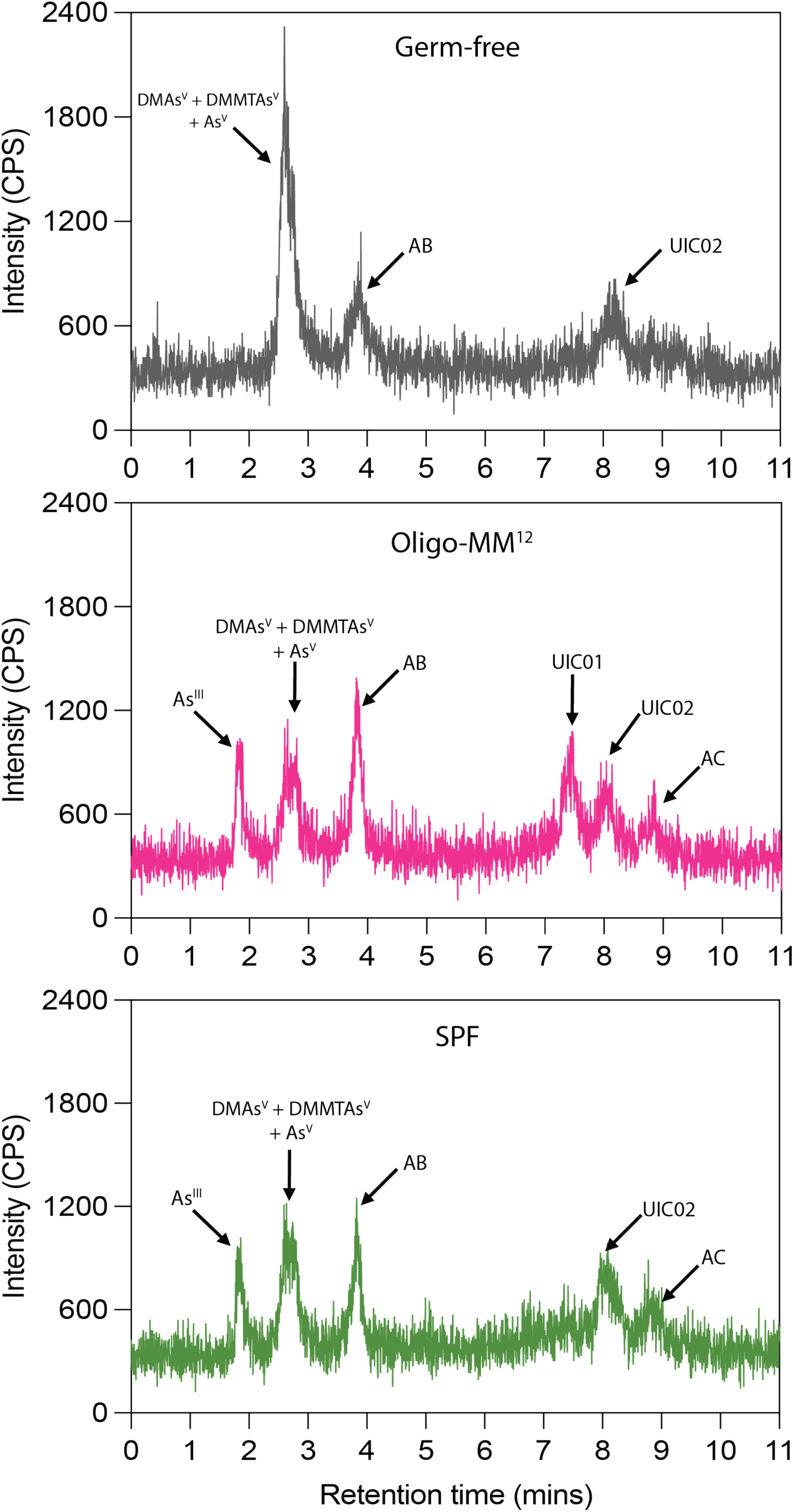
Cation exchange (CATEX) chromatogram of germ-free (GF) caecal content (top), Oligo-MM^12^ caecal content (middle) and conventional (SPF) (bottom). The Y-axis represents the scale for Intensity (CPS: counts per second) and the X-axis represents the retention time in minutes. The peaks were assigned to particular As species based upon a comparison with the chromatograms of the As species standards (Fig. S7). UIC01: Unidentified species 01 in CATEX; UIC02: Unidentified species 02 in CATEX.

### SECTION III

#### Faecal As concentrations

**Figure S9.**
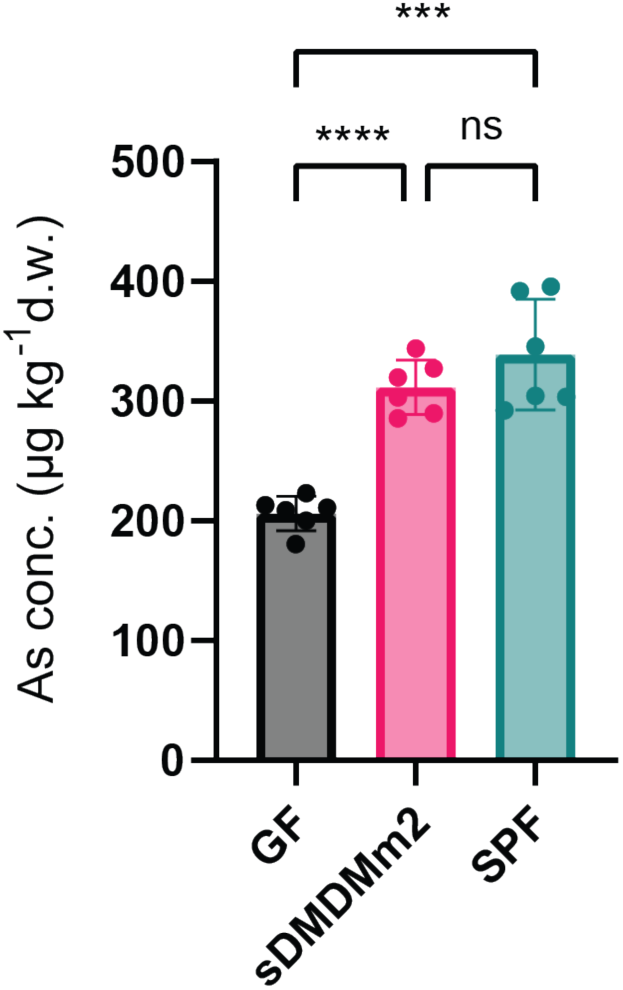
Total faecal As concentrations. Spot faecal samples of germ-free (GF, grey bars), gnotobiotic Oligo-MM12 (pink bars), and conventional (SPF, turquoise bars) mice following 6 weeks on AB-rich diet were studied. The samples were collected at a single time point after six weeks of feeding with the AB-rich diet. ****, adjusted p-value < 0.0001; ***, adjusted p-value = 0.0009; ns, non-significant, adjusted p-value = 0.3092; Brown-Forsythe and Welch ANOVA tests).

**Figure S10.**
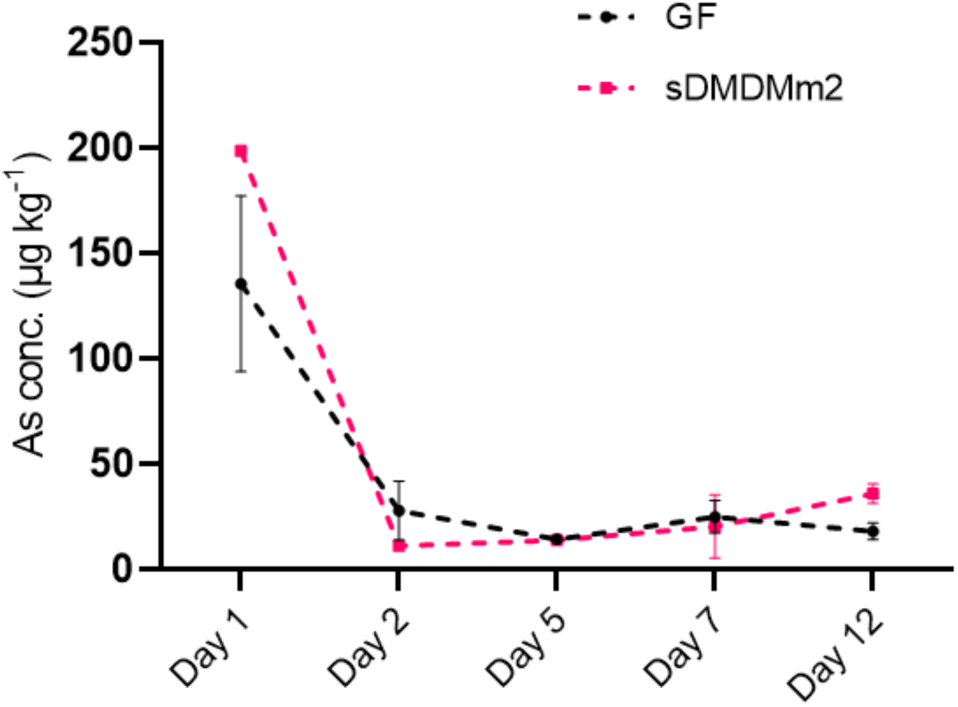
Time course of faecal As concentrations during 12 days in germ-free (GF) (black; n=2) and Oligo-MM^12^ (pink; n=2) mice following a switch from AB-rich rodent chow diet to low-As control diet.

## References

1. Tyson, J., The Determination of Arsenic Compounds: A Critical Review. ISRN Analytical Chemistry, 2013. 2013: p. 835371.

2. Matschullat, J., Arsenic in the geosphere — a review. Science of The Total Environment, 2000. 249(1): p. 297–312.

3. Raju, N.J., Arsenic in the geo-environment: A review of sources, geochemical processes, toxicity and removal technologies. Environmental Research, 2022. 203: p. 111782.

4. Naujokas, M.F., et al., The broad scope of health effects from chronic arsenic exposure: update on a worldwide public health problem. Environ Health Perspect, 2013. 121(3): p. 295–302.

5. Vahter, M., Health effects of early life exposure to arsenic. Basic Clin Pharmacol Toxicol, 2008. 102(2): p. 204–11.

6. Smith, A.H., et al., Evidence from Chile that arsenic in drinking water may increase mortality from pulmonary tuberculosis. Am J Epidemiol, 2011. 173(4): p. 414–20.

7. Islam, R., et al., Association between type 2 diabetes and chronic arsenic exposure in drinking water: a cross sectional study in Bangladesh. Environ Health, 2012. 11: p. 38.

8. Akter, K.F., et al., Arsenic speciation and toxicity in biological systems. Rev Environ Contam Toxicol, 2005. 184: p. 97–149.

9. Some metals and metallic compounds. IARC monographs on the evaluation of the carcinogenic risk of chemicals to humans, 1980. 23: p. 1–415.

10. Chi, L., et al., Individual susceptibility to arsenic-induced diseases: the role of host genetics, nutritional status, and the gut microbiome. Mammalian Genome, 2018. 29(1): p. 63–79.

11. Chi, H., et al., In vitro model insights into the role of human gut microbiota on arsenic bioaccessibility and its speciation in soils. Environ Pollut, 2020. 263(Pt A): p. 114580.

12. Yin, N., et al., Variability of arsenic bioaccessibility and metabolism in soils by human gut microbiota using different in vitro methods combined with SHIME. Sci Total Environ, 2016. 566**-**567: p. 1670-1677.

13. Calatayud, M., et al., Salivary and Gut Microbiomes Play a Significant Role in in Vitro Oral Bioaccessibility, Biotransformation, and Intestinal Absorption of Arsenic from Food. Environ Sci Technol, 2018. 52(24): p. 14422–14435.

14. Li, M.-Y., et al., Antibiotic exposure decreases soil arsenic oral bioavailability in mice by disrupting ileal microbiota and metabolic profile. Environment International, 2021. 151: p. 106444.

15. Coryell, M., et al., The gut microbiome is required for full protection against acute arsenic toxicity in mouse models. Nature Communications, 2018. 9(1): p. 5424.

16. Chi, L., et al., Gut microbiome disruption altered the biotransformation and liver toxicity of arsenic in mice. Arch Toxicol, 2019. 93(1): p. 25–35.

17. Song, D., et al., Gut microbiota promote biotransformation and bioaccumulation of arsenic in tilapia. Environmental Pollution, 2022. 305: p. 119321.

18. Van de Wiele, T., et al., Arsenic metabolism by human gut microbiota upon in vitro digestion of contaminated soils. Environ Health Perspect, 2010. 118(7): p. 1004–9.

19. Yin, N., et al., Interindividual variability of soil arsenic metabolism by human gut microbiota using SHIME model. Chemosphere, 2017. 184: p. 460–466.

20. Yin, N., et al., In Vitro Method To Assess Soil Arsenic Metabolism by Human Gut Microbiota: Arsenic Speciation and Distribution. Environ Sci Technol, 2015. 49(17): p. 10675–81.

21. Pinyayev, T.S., et al., Preabsorptive metabolism of sodium arsenate by anaerobic microbiota of mouse cecum forms a variety of methylated and thiolated arsenicals. Chem Res Toxicol, 2011. 24(4): p. 475–7.

22. Rowland, I.R. and M.J. Davies, In vitro metabolism of inorganic arsenic by the gastro-intestinal microflora of the rat. J Appl Toxicol, 1981. 1(5): p. 278–83.

23. Hall, L.L., et al., In VitroMethylation of Inorganic Arsenic in Mouse Intestinal Cecum. Toxicology and Applied Pharmacology, 1997. 147(1): p. 101–109.

24. Kubachka, K.M., et al., In vitro biotransformation of dimethylarsinic acid and trimethylarsine oxide by anaerobic microflora of mouse cecum analyzed by HPLC-ICP-MS and HPLC-ESI-MS. Journal of Analytical Atomic Spectrometry, 2009. 24(8).

25. Naranmandura, H., et al., Comparative toxicity of arsenic metabolites in human bladder cancer EJ-1 cells. Chem Res Toxicol, 2011. 24(9): p. 1586–96.

26. Dopp, E., et al., Subcellular Distribution of Inorganic and Methylated Arsenic Compounds in Human Urothelial Cells and Human Hepatocytes. Drug Metabolism and Disposition, 2008. 36(5): p. 971–979.

27. Moe, B., et al., Comparative cytotoxicity of fourteen trivalent and pentavalent arsenic species determined using real-time cell sensing. Journal of Environmental Sciences, 2016. 49.

28. Chen, F., et al., Sub-chronic low-dose arsenic in rice exposure induces gut microbiome perturbations in mice. Ecotoxicol Environ Saf, 2021. 227: p. 112934.

29. Gokulan, K., et al., Exposure to Arsenite in CD-1 Mice during Juvenile and Adult Stages: Effects on Intestinal Microbiota and Gut-Associated Immune Status. mBio, 2018. 9(4).

30. Harrington, C.F., E.I. Brima, and R.O. Jenkins, Biotransformation of arsenobetaine by microorganisms from the human gastrointestinal tract. Chemical Speciation & Bioavailability, 2008. 20(3): p. 173–180.

31. Fu, Y., et al., Arsenic speciation and bioaccessibility in raw and cooked seafood: Influence of seafood species and gut microbiota. Environmental Pollution, 2021. 280: p. 116958.

32. Popowich, A., Q. Zhang, and X.C. Le, Arsenobetaine: the ongoing mystery. National Science Review, 2016. 3(4): p. 451–458.

33. Luvonga, C., et al., Organoarsenicals in Seafood: Occurrence, Dietary Exposure, Toxicity, and Risk Assessment Considerations - A Review. J Agric Food Chem, 2020. 68(4): p. 943–960.

34. Taylor, V., et al., Human exposure to organic arsenic species from seafood. Sci Total Environ, 2017. 580: p. 266–282.

35. Cullen, W.R. and K.J. Reimer, Arsenic speciation in the environment. Chemical Reviews, 1989. 89(4): p. 713–764.

36. Dietary exposure to inorganic arsenic in the European population. EFSA Journal, 2014. 12(3).

37. Adams, M.A., Guidance document for arsenic in shellfish. 1993.

38. Jara, E.A. and C.K. Winter, Dietary exposure to total and inorganic arsenic in the United States, 2006–2008. International Journal of Food Contamination, 2014. 1(1): p. 3.

39. Borak, J. and H.D. Hosgood, Seafood arsenic: implications for human risk assessment. Regul Toxicol Pharmacol, 2007. 47(2): p. 204–12.

40. Schoof, R.A., et al., A market basket survey of inorganic arsenic in food. Food Chem Toxicol, 1999. 37(8): p. 839–46.

41. Yamauchi, H., et al., Intake of different chemical species of dietary arsenic by the Japanese, and their blood and urinary arsenic levels. Applied organometallic chemistry, 1992. 6(4): p. 383–388.

42. Kaise, T., S. Watanabe, and K. Itoh, The acute toxicity of arsenobetaine. Chemosphere, 1985. 14(9): p. 1327–1332.

43. Vahter, M., E. Marafante, and L. Dencker, Metabolism of arsenobetaine in mice, rats and rabbits. Science of The Total Environment, 1983. 30: p. 197–211.

44. Tam, G.K., et al., Excretion of a single oral dose of fish-arsenic in man. Bull Environ Contam Toxicol, 1982. 28(6): p. 669–73.

45. Freeman, H.C., et al., Clearance of arsenic ingested by man from arsenic contaminated fish. Bull Environ Contam Toxicol, 1979. 22(1-2): p. 224–9.

46. Yamauchi, H. and Y. Yamamura, Metabolism and excretion of orally ingested trimethylarsenic in man. Bull Environ Contam Toxicol, 1984. 32(6): p. 682–7.

47. Brown, R.M., et al., Human Metabolism of Arsenobetaine Ingested with Fish. Human & Experimental Toxicology, 1990. 9(1): p. 41–46.

48. Molin, M., et al., Humans seem to produce arsenobetaine and dimethylarsinate after a bolus dose of seafood. Environmental Research, 2012. 112: p. 28–39.

49. Newcombe, C., et al., Accumulation or production of arsenobetaine in humans? Journal of Environmental Monitoring, 2010. 12(4): p. 832–837.

50. Lai, V.W.M., et al., Arsenic speciation in human urine: are we all the same? Toxicology and Applied Pharmacology, 2004. 198(3): p. 297–306.

51. Ye, Z., et al., Biodegradation of arsenobetaine to inorganic arsenic regulated by specific microorganisms and metabolites in mice. Toxicology, 2022. 475: p. 153238.

52. Zhang, J., et al., Significant Biotransformation of Arsenobetaine into Inorganic Arsenic in Mice. Toxics, 2023. 11(2).

53. Brugiroux, S., et al., Genome-guided design of a defined mouse microbiota that confers colonization resistance against Salmonella enterica serovar Typhimurium. Nat Microbiol, 2016. 2: p. 16215.

54. Authority, E.F.S., Opinion of the Scientific Panel on contaminants in the food chain [CONTAM] related to Arsenic as undesirable substance in animal feed. EFSA Journal, 2005. 3(3): p. 180.

55. Yoshida, K., et al., Urinary excretion of arsenic metabolites after long-term oral administration of various arsenic compounds to rats. J Toxicol Environ Health A, 1998. 54(3): p. 179–92.

56. Yoshida, K., et al., Metabolites of arsenobetaine in rats: does decomposition of arsenobetaine occur in mammals? Applied Organometallic Chemistry, 2001. 15(4): p. 271–276.

57. Pandey, N. and R. Bhatt, Arsenic resistance and accumulation by two bacteria isolated from a natural arsenic contaminated site. J Basic Microbiol, 2015. 55(11): p. 1275–86.

58. Takeuchi, M., et al., Arsenic resistance and removal by marine and non-marine bacteria. J Biotechnol, 2007. 127(3): p. 434–42.

59. Naranmandura, H., K. Ibata, and K.T. Suzuki, Toxicity of dimethylmonothioarsinic acid toward human epidermoid carcinoma A431 cells. Chem Res Toxicol, 2007. 20(8): p. 1120–5.

60. Jenkins, R.O., et al., Bacterial degradation of arsenobetaine via dimethylarsinoylacetate. Arch Microbiol, 2003. 180(2): p. 142–50.

61. Chen, J., et al., Organoarsenical compounds: Occurrence, toxicology and biotransformation. Critical Reviews in Environmental Science and Technology, 2020. 50(3): p. 217–243.

62. Khokiattiwong, S., et al., Dimethylarsinoylacetate from microbial demethylation of arsenobetaine in seawater. Applied Organometallic Chemistry, 2001. 15(6): p. 481–489.

63. Hanaoka, K.i., et al., Arsenobetaine-decomposing Ability of Marine Microorganisms Occurring in Particles Collected at Depths of 1100 and 3500 Meters. Applied Organometallic Chemistry, 1997. 11(4): p. 265–271.

64. SS, D.C.R., et al., Arsenic thiolation and the role of sulfate-reducing bacteria from the human intestinal tract. Environ Health Perspect, 2014. 122(8): p. 817–22.

65. Carbonero, F., et al., Microbial pathways in colonic sulfur metabolism and links with health and disease. Front Physiol, 2012. 3: p. 448.

66. Naranmandura, H., et al., Comparative Toxicity of Arsenic Metabolites in Human Bladder Cancer EJ-1 Cells. Chemical Research in Toxicology, 2011. 24(9): p. 1586–1596.

67. Wang, P., et al., Role of human gut bacteria in arsenic biosorption and biotransformation. Environment International, 2022. 165: p. 107314.

68. McDermott, T.R., J.F. Stolz, and R.S. Oremland, Arsenic and the gastrointestinal tract microbiome. Environ Microbiol Rep, 2020. 12(2): p. 136–159.

69. Wang, Q.Q., D.J. Thomas, and H. Naranmandura, Importance of being thiomethylated: formation, fate, and effects of methylated thioarsenicals. Chem Res Toxicol, 2015. 28(3): p. 281–9.

70. Devesa, V., et al., Effect of Cooking Temperatures on Chemical Changes in Species of Organic Arsenic in Seafood. Journal of Agricultural and Food Chemistry, 2001. 49(5): p. 2272–2276.

71. Dahl, L., et al., Stability of arsenic compounds in seafood samples during processing and storage by freezing. Food Chemistry, 2010. 123: p. 720–727.

72. Laird, B.D., et al., Gastrointestinal Microbes Increase Arsenic Bioaccessibility of Ingested Mine Tailings Using the Simulator of the Human Intestinal Microbial Ecosystem. Environmental Science & Technology, 2007. 41(15): p. 5542–5547.

73. Yin, N., et al., Predictive capabilities of in vitro colon bioaccessibility for estimating in vivo relative bioavailability of arsenic from contaminated soils: Arsenic speciation and gut microbiota considerations. Sci Total Environ, 2022. 818: p. 151804.

74. Wostmann, B.S., et al., Dietary intake, energy metabolism, and excretory losses of adult male germfree Wistar rats. Laboratory animal science, 1983. 33(1): p. 46–50.

75. Bäckhed, F., et al., The gut microbiota as an environmental factor that regulates fat storage. Proc Natl Acad Sci U S A, 2004. 101(44): p. 15718–23.

76. Gregus, Z., Á. Gyurasics, and I. Csanaky, Biliary and Urinary Excretion of Inorganic Arsenic: Monomethylarsonous Acid as a Major Biliary Metabolite in Rats. Toxicological Sciences, 2000. 56(1): p. 18–25.

77. Suzuki, K.T., et al., Glutathione-conjugated arsenics in the potential hepato-enteric circulation in rats. Chem Res Toxicol, 2001. 14(12): p. 1604–11.

78. Hughes, M.F., et al., Accumulation and metabolism of arsenic in mice after repeated oral administration of arsenate. Toxicology and Applied Pharmacology, 2003. 191(3): p. 202–210.

79. Gregus, Z. and C.D. Klaassen, Disposition of metals in rats: A comparative study of fecal, urinary, and biliary excretion and tissue distribution of eighteen metals. Toxicology and Applied Pharmacology, 1986. 85(1): p. 24–38.

80. Hackethal, C., et al., Total arsenic and water-soluble arsenic species in foods of the first German total diet study (BfR MEAL Study). Food Chem, 2021. 346: p. 128913.

81. Grinham, A., et al., Baseline arsenic levels in marine and terrestrial resources from a pristine environment: Isabel Island, Solomon Islands. Marine Pollution Bulletin, 2014. 88(1): p. 354–360.

82. Yilmaz, B., et al., Long-term evolution and short-term adaptation of microbiota strains and sub-strains in mice. Cell Host Microbe, 2021. 29(4): p. 650–663.e9.

83. Chen, B., et al., Identification of Methylated Dithioarsenicals in the Urine of Rats Fed with Sodium Arsenite. Chemical Research in Toxicology, 2016. 29(9): p. 1480–1487.

84. Au- Lee, H., et al., Preparation of DMMTAV and DMDTAV Using DMAV for Environmental Applications: Synthesis, Purification, and Confirmation. JoVE, 2018(133): p. e56603.

85. McKnight-Whitford, A., et al., New Method and Detection of High Concentrations of Monomethylarsonous Acid Detected in Contaminated Groundwater. Environmental Science & Technology, 2010. 44(15): p. 5875–5880.

86. Chávez-Capilla, T., et al., Bioaccessibility and degradation of naturally occurring arsenic species from food in the human gastrointestinal tract. Food Chemistry, 2016. 212: p. 189–197.

87. Francesconi, K.A. and J.S. Edmonds, Arsenic and Marine Organisms, in Advances in Inorganic Chemistry, A.G. Sykes, Editor. 1996, Academic Press. p. 147–189.

88. Christakopoulos, A., et al., Cellular metabolism of arsenocholine. J Appl Toxicol, 1988. 8(2): p. 119–27.

